# A cyanobacterial adenine prenyltransferase enables longer-chain N6 prenylation

**DOI:** 10.64898/2026.04.03.716432

**Authors:** Kotoha Ichikawa, Kentaro Tamura, Koki Fujitani, Taichi Chisuga, Ryunosuke Takeda, Tomoki Sato, Seiichiro Hayashi, Atsuji Kodama, Koichi Kato, Shinji Miura, Shogo Nakano, Sohei Ito, Daisuke Fujinami

## Abstract

Adenine, a ubiquitous nucleobase found in biomolecules such as nucleic acids, cofactors, and signaling molecules, mediates diverse molecular interactions. Here, we identify TvAPT, an adenine prenyltransferase from the cyanobacterium *Trichormus variabilis* NIES-23. TvAPT efficiently catalyzes the unprecedented N6-prenylation of adenine-containing substrates using extended C10 and C15 prenyl donors, whereas canonical adenine prenyltransferases are generally limited to C5-dimethylallylation. X-ray structural analyses and protein engineering revealed that an enlarged prenyl-binding pocket enables the accommodation of extended prenyl donors, providing a structural basis for rationally tuning prenyl-donor preference. Together, these findings establish TvAPT as an adenine prenyltransferase that accepts a range of adenine-containing substrates, with a preference for nucleoside 5’-monophosphates. Beyond expanding the known catalytic scope of adenine prenyltransferases, TvAPT provides access to nucleotide derivatives with enhanced membrane permeability and analogues of plant signaling molecules.

## Introduction

Adenine is a ubiquitous purine nucleobase that is essential for life and serves as one of the canonical building blocks of nucleic acids. In DNA and RNA, it contributes to higher-order structure through base pairing^1^. Beyond nucleic acids, adenine is incorporated into a wide range of biomolecules, including ATP, cAMP, NAD, FAD, coenzyme A and SAM ^2,3^. In many of these molecules, the adenine moiety functions primarily as a molecular recognition element via hydrogen-bonding and π-interactions, rather than acting as the principal chemically reactive center^4^. Because of this versatile recognition capacity, adenine-containing scaffolds are prevalent in diverse natural products and therapeutic agents, exerting potent bioactivities in nucleic acid synthesis, enzyme function, and ribosomal translation^5–8^. However, the poor membrane permeability of adenine nucleotides restricts their direct intracellular utility ^9^. Instead, when exported into the extracellular space via transporters and channels, adenine nucleotides act as signaling molecules after being dephosphorylated to adenosine.

Modifications at the N6 nitrogen of adenine are widespread in biological systems. In RNA, modifications such as N6-methyladenosine (m^6^A), N6-threonylcarbamoyladenosine (t6A), and N6-isopentenyladenosine (i^6^A) regulate critical processes ranging from mRNA splicing^10^ to ribosomal translation fidelity^11,12^ (Supplementary Fig. 1). Specifically, i^6^A is generated by isopentenyltransferases (IPTs) through the transfer of a C5-dimethylallyl group^13^, and this modification has also been linked to tRNA membrane association ^14^. i^6^A can be further enzymatically hypermodified to generate derivatives such as 2-methylthio-N6-(cis-hydroxyisopentenyl)adenosine (ms^2^io^6^A)^13^. Beyond RNA, N6-prenylation plays a crucial role in cellular signaling as a modification of monomeric nucleotides. In plants and *Agrobacterium*, adenylate IPTs catalyze the C5-dimethylallylation of AMP, which serves as the initial step in the biosynthesis of cytokinins, phytohormones that are central to plant growth and stress responses^15–17^ (Fig. 1A).

Long-chain isoprenoid groups are widely utilized in protein post-translational modifications, where longer chain lengths enhance membrane anchoring and modulate subcellular localization^18,19^. Similarly, conjugation of nucleic acids with long-chain lipids, particularly acyl chains or cholesterol, has been exploited to enhance the cellular uptake and tissue-selective delivery of oligonucleotide therapeutics^20,21^ and to construct nucleic acid assemblies on lipid membranes^22^. Although long-chain prenylation has been reported for uridine derivatives, including S-geranylation^23^ and 5’-O-geranylation^24^ (Supplementary Fig. 1), naturally occurring N6-prenylation of adenine has, to our knowledge, been limited to C5-dimethylallylation ^25^. By analogy with protein prenylation, we hypothesized that extending the N6-prenylation of adenine beyond the C5-dimethylallyl group would expand the chemical space of nucleobases and give rise to novel biological functions.

Here, we characterize TvAPT, an adenine prenyltransferase from the cyanobacterium *Trichormus variabilis* NIES-23. TvAPT catalyzes the unprecedented N6-prenylation of adenine-containing substrates using extended prenyl donors, with the particularly efficient transfer of C10 and C15 groups and weaker but detectable activity toward C20 donors (Fig. 1B). Structural and mutational analyses revealed that an enlarged prenyl-binding pocket underlies this expanded donor scope and identified a key residue that controls the prenyl-donor preference. TvAPT also accepts a range of adenine-containing substrates, with a preference for nucleoside 5’-monophosphates. Using TvAPT, we synthesized membrane-permeable fluorescent AMP derivatives and analogues of plant signaling molecules. Overall, these results establish TvAPT as a biocatalytic tool for adenine diversification and provide a foundation for future applications in biomolecular engineering.

**Figure 1.**
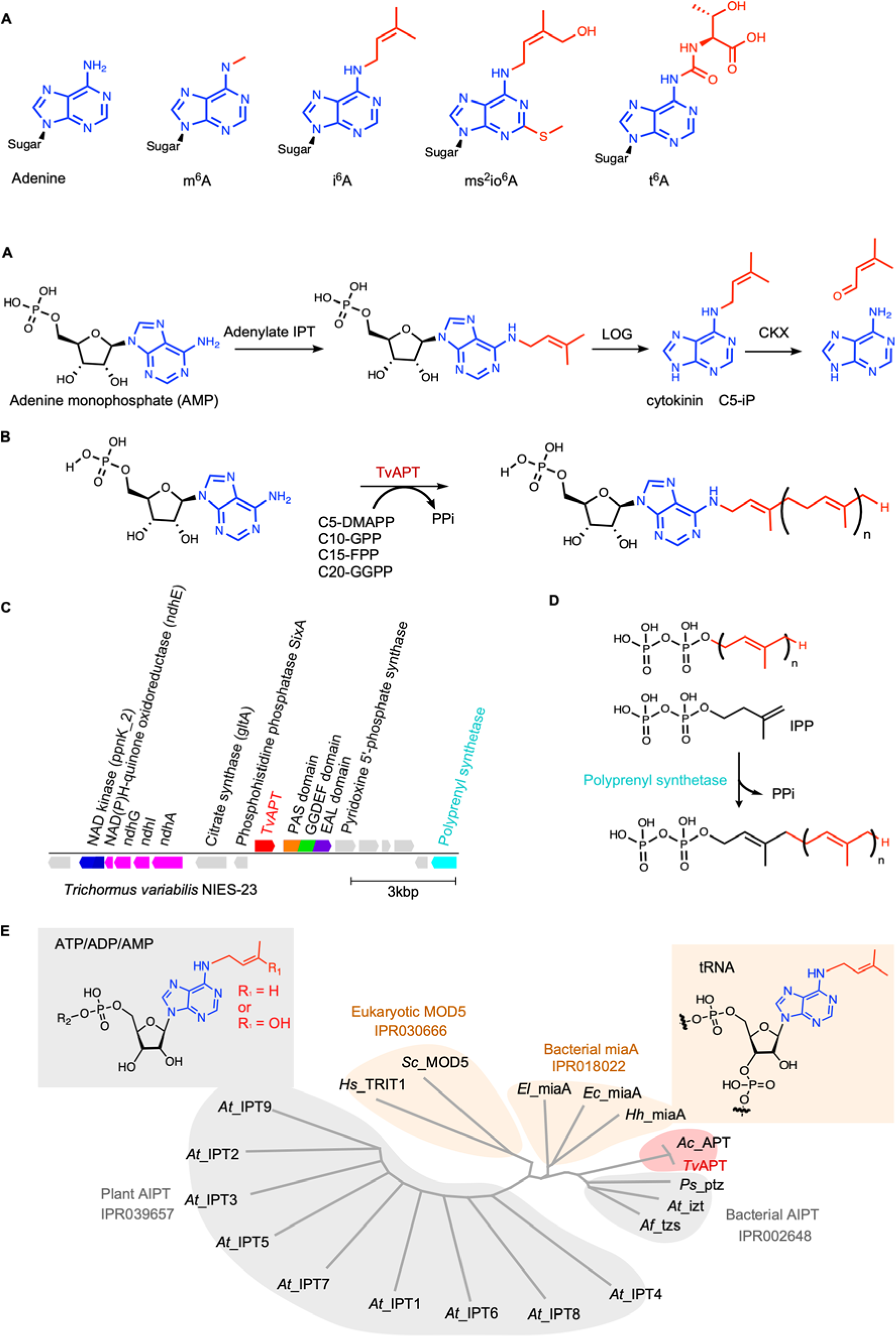
Discovery and evolutionary context of the cyanobacterial adenine prenyltransferase TvAPT. (A) Biosynthesis and degradation pathways of the cytokinin isopentenyladenine (C5-iP). The adenine moiety is shown in blue, and the modifying groups are shown in red. (B) Reaction scheme for the TvAPT-catalyzed N6-prenylation of adenine-containing substrates using C5-dimethylallyl diphosphate (DMAPP), C10-geranyl diphosphate (GPP), C15-farnesyl diphosphate (FPP), or C20-geranylgeranyl diphosphate (GGPP) as prenyl donors. (C) Genomic organization of the region surrounding the TvAPT locus. (D) Proposed prenyl-chain elongation mediated by a putative polyprenyl synthase. (E) Phylogenetic analysis of adenine prenyltransferases. Clades containing nucleotide- and tRNA-prenylating enzymes are highlighted in gray and orange, respectively, with cyanobacterial APTs indicated in red.

## Results

### Identification of TvAPT through a genome-mining approach

To discover adenine prenyltransferases with potentially non-canonical functions, we constructed a sequence similarity network (SSN) using the EFI-EST tool ^26^. We built the network with two protein families: adenylate dimethylallyltransferases involved in bacterial cytokinin biosynthesis (InterPro: IPR002648) and dimethylallyltransferases mediating tRNA modification and plant cytokinin biosynthesis (IPR039657) (Supplementary Fig. 2). The latter includes both eukaryotic (IPR030666) and predominantly bacterial (IPR018022) tRNA isopentenyltransferases (Fig. 1E). By analyzing genomic contexts via EFI-GNT, we prioritized putative adenine prenyltransferase genes lacking genomic proximity to canonical tRNA-modifying enzymes or the cytokinin riboside 5’-monophosphate phosphoribohydrolase, LOG (IPR005269) (Fig. 1A and Supplementary Fig. 3). This strategy highlighted a candidate locus in the cyanobacterium *Trichormus variabilis* NIES-23 encoding a putative adenine prenyltransferase, which we named TvAPT (UniProt: A0A1Z4KQD0). Notably, the TvAPT gene co-clusters with genes encoding an NAD kinase, NADH-quinone oxidoreductase subunits (ndh), a GGDEF–EAL domain-containing protein, and a polyprenyl synthetase-like protein (Fig. 1C). Although TvAPT is identical in sequence to a previously reported adenylate dimethylallyltransferase from Nostoc sp. PCC 7120 ^27^, this specific gene cluster organization is restricted to a narrow group of cyanobacteria, primarily *Anabaena*, *Nostoc*, and *Trichormus*^28,29^ (Supplementary Figs. 2 and 19). Phylogenetically, TvAPT clusters more closely with bacterial AIPTs than with canonical tRNA isopentenyltransferases (Fig. 1E).

Several cyanobacteria are known to produce the cytokinin isopentenyladenine (C5-iP), but we could not identify a canonical LOG homologue in the *Trichormus variabilis* NIES-23 genome^27,29–32^. Nevertheless, we found a gene annotated as cytochrome d ubiquinol oxidase subunit II (UniProt: A0A1Z4KQ66) in a region distant from the TvAPT locus. This protein belongs to the broader LOG family (IPR031100), which includes the pyrimidine/purine nucleotide 5’-monophosphate nucleosidase YgdH. Although its precise cellular function remains undefined, it may contribute to cyanobacterial cytokinin biosynthesis^27^. Furthermore, we identified a CKX-like protein (UniProt: A0A1Z4KQP8) in the genome, suggesting the presence of a cyanobacterial cytokinin degradation pathway (Fig. 1A).

### An enlarged binding pocket enables long-chain adenine prenylation by TvAPT

The genomic co-localization of TvAPT with a polyprenyl synthase, an enzyme predicted to generate prenyl chains longer than C5^33^, led us to hypothesize that TvAPT might accept prenyl donors beyond dimethylallyl diphosphate (DMAPP) ^15,25^. To test this, we compared the prenyl-donor preferences of recombinant TvAPT and the *Agrobacterium fabrum* AIPT (tzs) *in vitro*. Since NolPT1, an identical homologue of TvAPT from *Nostoc* sp. PCC 7120, was previously shown to utilize AMP as a substrate for C5-dimethylallylation^27^, we selected AMP as the prenyl acceptor. We incubated TvAPT or tzs with AMP and MgCl_2_ in the presence of an equimolar mixture of prenyl diphosphates: C5-DMAPP, C10-geranyl diphosphate (GPP), C15-farnesyl diphosphate (FPP), and C20-geranylgeranyl diphosphate (GGPP). HPLC analysis revealed broad donor promiscuity for TvAPT, which preferentially utilized C10-GPP (77.1% relative activity) and C15-FPP (13.9%) alongside C5-DMAPP (1.8%) (Fig. 2A and Supplementary Fig. 4). Although C20-geranylgeranylated AMP was not detected in this competition assay, incubating TvAPT with GGPP as the sole prenyl donor yielded a detectable, albeit minor, amount of the C20-prenylated product (Supplementary Fig. 4). By contrast, under identical conditions, tzs produced only the C5-dimethylallylated AMP. Together, these results demonstrate that TvAPT possesses an unprecedented capacity to accept prenyl donors ranging from C5 to C20.

To elucidate the structural basis for this donor promiscuity, we used Boltz-2 ^34^ to predict the structures of TvAPT complexed with AMP and C15-FPP, and tzs complexed with AMP and DMAPP (Fig. 2B and Supplementary Fig. 5). Using PyVOL ^35^, subsequent volume estimations of the prenyl-donor binding pockets yielded values of 351 Å^3^ for TvAPT and 122 Å^3^ for tzs, confirming a substantially enlarged cavity in TvAPT. Structural comparisons highlighted Ala142 in TvAPT as a critical residue contributing to this expanded pocket architecture. To directly assess its role, we generated the TvAPT A142M mutant and the reciprocal tzs M142A mutant. In prenyl-competition assays, the TvAPT A142M mutant exhibited a clear shift in donor preference from C10-GPP back to C5-DMAPP, underscoring the pivotal role of Ala142 as a key determinant of prenyl-donor selectivity (Fig. 2A). Conversely, while the tzs M142A mutant gained a marginal ability to produce geranylated AMP, its primary preference did not shift toward C10-GPP. Overlay of the overall structures revealed subtle shifts in helices α4, α7, and α8 in TvAPT, consistent with a slightly more open prenyl-binding pocket (Fig. 2C).

**Figure 2.**
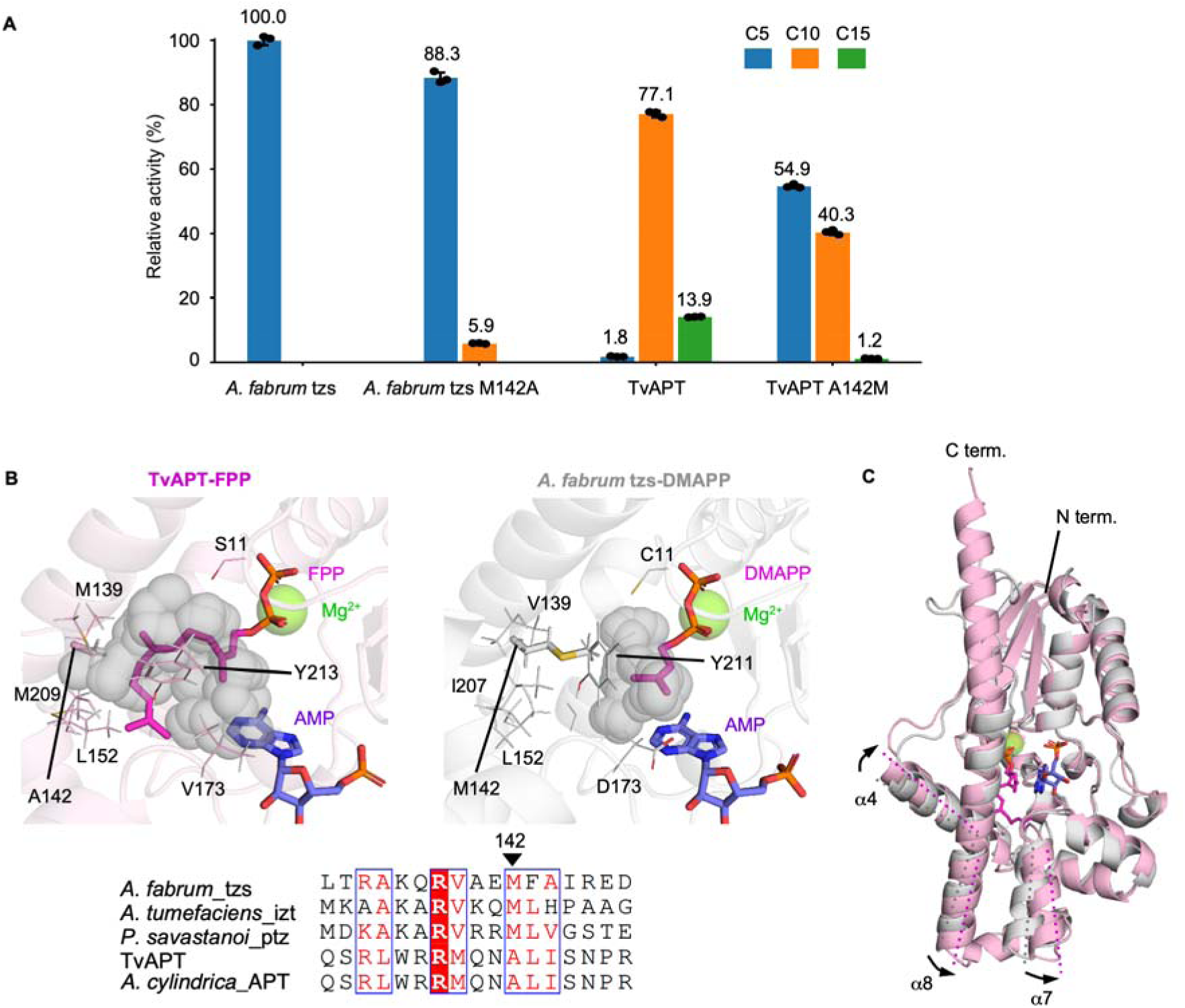
Comparison of prenyl-donor specificities between TvAPT and tzs from *A. fabrum*. (A) Prenyl-donor competition assay using an equimolar mixture of C5, C10, C15, and C20 prenyl donors. Relative activities were calculated from the integrated HPLC peak areas of the corresponding prenylated products. The corresponding HPLC chromatograms are shown in Supplementary Fig. 4. Data are presented as the mean relative activity ± s.d. from three independent experiments. Mean values are shown above the bars. (B) Predicted prenyl-binding sites of TvAPT (pink, left) and tzs (gray, right). The sequence alignment below shows residues surrounding position 142. (C) Structural overlay of TvAPT and tzs. Differences in α-helix positions are indicated by black arrows.

### Computational design enables crystallographic determination of product- and substrate-analogue-bound TvAPT_20per structures

Initial crystallization trials of wild-type TvAPT were hampered by limited protein stability and poor crystallizability. To overcome these challenges, we implemented a computational protein-design strategy. Using ESM-scan^36^ to predict site-specific effects on stability, we selected 25 residues (approximately 10% of the 244-residue sequence) for redesign while retaining residues predicted to be important for catalysis, including those involved in acceptor AMP binding, prenyl diphosphate donor binding, and metal-ion coordination. These positions were substituted with amino acids proposed by LigandMPNN^37^, a deep learning-based protein design method that incorporates local sequence context. This approach generated a total of 20 redesigned sequence variants, which were subsequently integrated into a single sequence via ancestral sequence reconstitution, yielding a variant we termed TvAPT_20per (Fig. 3A).

Although TvAPT_20per exhibited lower geranyltransferase activity than the wild-type enzyme, it retained the ability to modify AMP (Fig. 3C). We therefore determined the steady-state kinetic parameters of wild-type TvAPT and TvAPT_20per, using AMP as the variable substrate (Fig. 3C and Supplementary Fig. 6). Because the GPP concentration was fixed at 500 μM to minimize complications arising from the limited solubility of the reaction product, the resulting values represent apparent kinetic parameters. The *K*_□,app_ values for AMP were similar, at 7.6 μM for wild-type TvAPT and 13.7 μM for TvAPT_20per. By contrast, TvAPT_20per showed a pronounced reduction in catalytic turnover, with *k*_cat,app_ values of 1.64 min□^1^ for the wild-type enzyme and 0.05 min□^1^ for TvAPT_20per, corresponding to an approximately 33-fold decrease. Thus, the redesign affected the catalytic turnover much more strongly than the AMP-dependent Michaelis constant under the assay conditions. The pronounced reduction in *k*_cat,app_ may reflect perturbations of conformational changes or other dynamic processes required for efficient catalysis.

Because TvAPT_20per still formed amorphous precipitates during prolonged incubation at room temperature, we conducted additive screening and identified sodium sulfate as a potent solubility enhancer. Ultimately, co-crystallization in the presence of 300 mM Na SO, 1 mM GPP, and 5 mM MgCl yielded diffraction-quality crystals, allowing us to determine the structure at 1.7 Å resolution (PDB: 24WM). Within the crystal structure, three sulfate ions were bound to solvent-exposed, positively charged regions on the protein surface, consistent with their observed stabilizing effect (Fig. 3B). At the active site, we detected electron density consistent with a low-occupancy geranylated AMP molecule, despite the absence of exogenous AMP in the crystallization buffer (Fig. 3D). This ligand likely originated from the *E. coli* expression host and co-purified tightly with TvAPT. The electron density defined the position of the geranyl C1 atom and was consistent with covalent attachment to the N6 position of AMP in the forward orientation. The identity and connectivity of the N6-geranylated product were independently supported by NMR and high-resolution MS analyses of the purified *in vitro* reaction product (Fig. 3E and Supplementary Figs. 7 and 8).

To investigate substrate recognition before prenyl transfer, we next determined a substrate-analogue-bound structure. Co-crystallization with 300 mM Na_2_SO_4_, 2 mM GSPP (a sulfur-containing analogue of GPP in which one of the bridging oxygen atoms is replaced by sulfur), 2 mM AMP, and 5 mM MgCl_2_ yielded diffraction-quality crystals. The structure was determined at 1.85 Å resolution (Fig. 3F; PDB: 27DP). In the TvAPT_20per structure, the 2’-hydroxyl group of AMP forms a hydrogen bond with Thr171, suggesting that this interaction contributes to substrate recognition. To assess the functional importance of the 2’-hydroxyl group in the wild-type enzyme, we performed a steady-state kinetic analysis using 2’-deoxyadenosine monophosphate (dAMP), which lacks this group, as the acceptor substrate (Supplementary Fig. 6). The apparent Michaelis constant increased from 7.6 μM for AMP to 51.5 μM for dAMP, corresponding to an approximately 6.8-fold increase, whereas the apparent turnover number only decreased from 1.64 to 0.90 min ¹. The preferential effect on *K*_□, app_ suggests that the 2’-hydroxyl group contributes primarily to productive substrate recognition, rather than catalytic turnover. These kinetic results obtained with wild-type TvAPT are consistent with the local AMP-recognition geometry observed in the TvAPT_20per structure and suggest that the engineered and wild-type enzymes share a similar mode of 2’-hydroxyl recognition.

The 5’-phosphate group of AMP is positioned to form hydrogen bonds with the N_ε_ atom of Arg34 and the side-chain hydroxyl groups of Ser100 and Ser102. The functional roles of these interactions are examined in the following subsection. To explore the structural differences between the substrate-analogue- and product-bound states, we compared the two TvAPT_20per structures. Their overall conformations were highly similar (Supplementary Fig. 9). Despite this overall similarity, AMP and geranylated AMP adopted distinct bound conformations. In the product-bound structure, the adenine ring of geranylated AMP was oriented toward the prenyl diphosphate-binding site (Fig. 3H). Asp33 is highly conserved among related adenine prenyltransferases and lies within hydrogen-bonding distance of the N6 atom of AMP in the substrate-analogue-bound structure (3.0 Å; Fig. 3H and Supplementary Fig. 10). This geometry is consistent with a possible role for Asp33 in orienting and polarizing the N6 amino group for nucleophilic attack on the C1 atom of the prenyl diphosphate donor (Supplementary Fig. 11). The N6 atom of AMP is positioned 3.6 Å from the C1 atom of GSPP, placing the two atoms in a geometry compatible with nucleophilic attack. Reactions catalyzed by related prenyltransferases have been proposed to proceed through either an S_N_1-like dissociative pathway, in which prenyl donor ionization and pyrophosphate departure precede nucleophilic attack, or a more concerted S_N_2-like pathway ^38,39^(Supplementary Fig. 11). The observed geometry is compatible with either mechanism, and the present structural and biochemical data do not distinguish between them.

To further probe possible catalytic intermediates by altering the chemical properties of the acceptor substrate, we performed the geranylation reaction using N6-methyl-AMP. HPLC analysis of the reaction mixture revealed two product peaks, which were assigned by MALDI-TOF mass spectrometry to N6-methyl-N6-geranyl-AMP and N6-geranyl-AMP (Supplementary Fig. 12), with an approximate UV peak-area ratio of 6:4. One possible interpretation is that geranyl transfer initially generates a cationic N6-methyl-N6-geranyl intermediate. Deprotonation of this intermediate would yield N6-methyl-N6-geranyl-AMP, whereas competing cleavage of the N6–CH_3_ bond through an N-dealkylation-like process could account for the formation of N6-geranyl-AMP.

Notably, no bound metal ion could be modeled in the active site of the substrate-analogue-bound structure (Fig. 3F), in contrast to the metal ions observed in crystal structures of related prenyltransferases^15,40,41^. Although MgCl_2_ was included in the crystallization solution, the absence of a resolvable metal ion may be attributable to chelation by citrate in the crystallization solution, dissociation during cryoprotection because MgCl_2_ was absent from the cryoprotectant solution, low metal occupancy, or insufficiently ordered coordination. To systematically evaluate the metal dependence of TvAPT catalysis, we prepared metal-depleted TvAPT via EDTA treatment and dialysis. This enzyme was then incubated with AMP and GPP in the presence of various divalent metal cations (Zn^2+^, Ni^2+^, Mn^2+^, Cu^2+^, Mg^2+^, Ca^2+^, and Co^2+^). No geranylation occurred in the absence of supplemented metal ions. Conversely, product formation was restored by all tested metal ions, although Zn^2+^ supported comparatively lower activity (Supplementary Fig. 13). To investigate whether catalytic activity correlated with Pearson hardness, an empirical indicator of a metal ion’s affinity for hard bases such as phosphate oxygens, we plotted relative activity against Pearson hardness; however, no clear trend was observed (Supplementary Fig. 13) ^42^. Because the measured activity reflects multiple factors, including metal-dependent substrate recognition, coordination geometry, and subsequent chemical steps, Pearson hardness alone may not fully account for the observed metal dependence. Consistent with this possibility, divalent metal ions in related prenyltransferases have been proposed not only to coordinate the prenyl diphosphate donor but also to facilitate donor ionization and stabilize the departing diphosphate group^43^. To explore whether the different metal ions primarily affected donor recognition or catalytic turnover, we performed preliminary steady-state kinetic measurements using GPP as the variable substrate in the presence of Mg^2+^ or Zn^2+^. However, sufficiently well-defined substrate-saturation curves could not be obtained over the experimentally accessible GPP concentration range, precluding reliable determinations of the apparent *K* and *k*cat values. The present data therefore do not allow us to determine whether the metal dependence of TvAPT arises primarily from effects on prenyl donor recognition, promotion of diphosphate departure, or a combination of these processes. Direct measurements of GPP binding under different metal conditions, for example by isothermal titration calorimetry, may help distinguish effects on donor binding from effects on subsequent catalytic steps.

**Figure 3.**
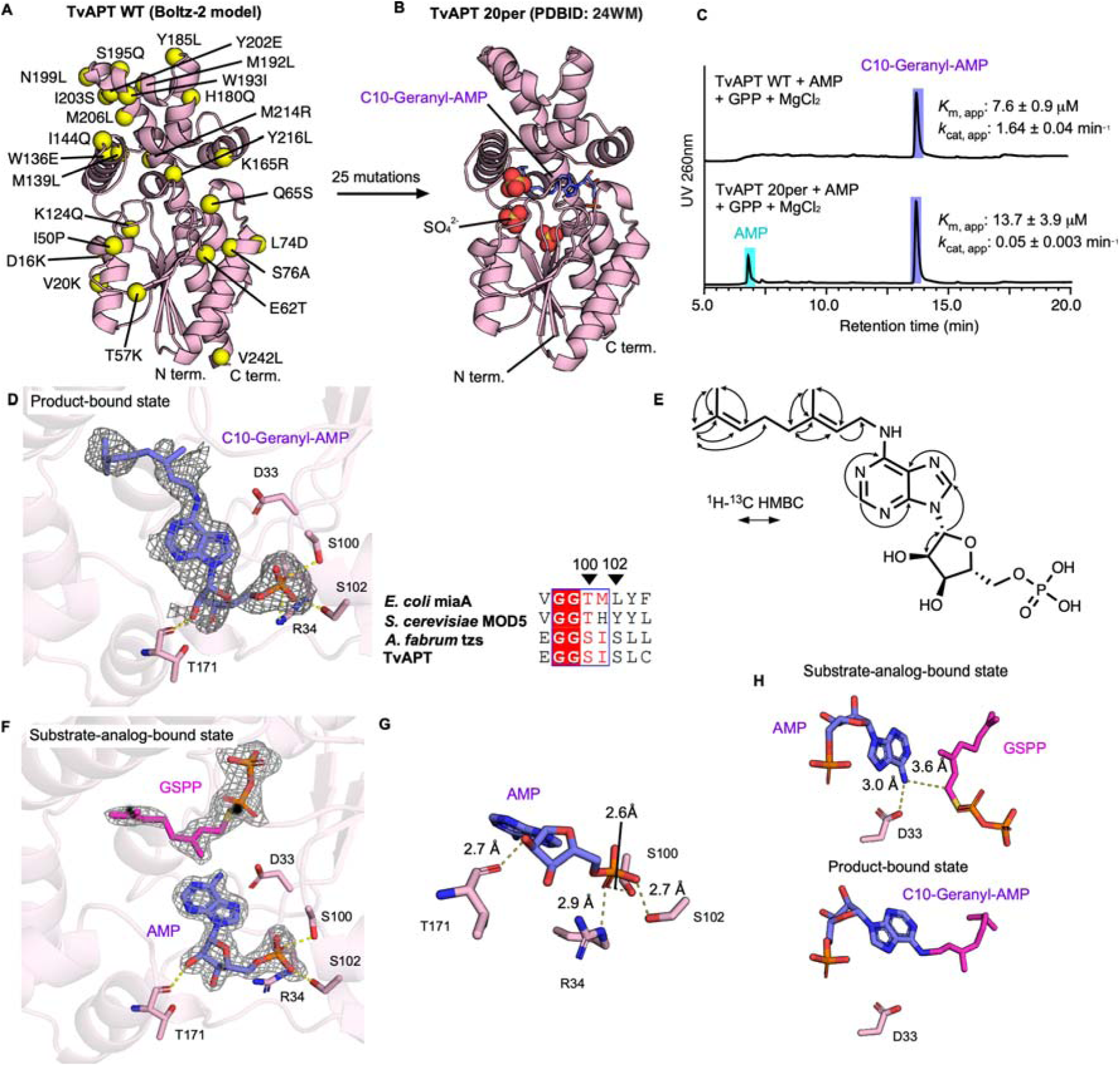
Crystallographic analysis of the engineered TvAPT variant. (A) The 25 computationally redesigned residues mapped onto the Boltz-2 structural model of TvAPT. (B) Crystal structure of geranyl-AMP-bound TvAPT_20per. Geranyl-AMP is shown as purple sticks, and the three bound sulfate ions are shown as spheres. (C) HPLC analysis of the *in vitro* prenyltransferase activities of wild-type TvAPT and TvAPT_20per. The apparent *K*_m_ and *k*_cat_ values determined by kinetic analysis in Supplementary Fig. 6 are shown alongside the corresponding chromatograms. (D) Close-up view of the active site of geranyl-AMP-bound TvAPT_20per. The F_o_–F_c_ omit electron density map for geranyl-AMP is shown as a gray mesh and contoured at 1.0 σ. The accompanying multiple sequence alignment (right) highlights the conserved residues corresponding to Ser100 and Ser102 in TvAPT, which are involved in 5’-monophosphate group recognition. (E) Chemical structure of N6-geranyl-AMP showing the key ^1^H–^13^C HMBC correlations. (F) Close-up view of the active site of substrate-analogue-bound TvAPT_20per. The F_o_–F_c_ omit electron-density maps are shown as gray meshes and contoured at 2.0 σ for GSPP and 3.0 σ for AMP. (G) Recognition of the 2’-hydroxyl and 5’-phosphate groups of AMP in the active site. AMP and the interacting residues are shown as sticks. Putative hydrogen bonds are shown as yellow dashed lines, with distances indicated in Å. (H) Comparison of the bound conformations of AMP and geranylated AMP. The distances between the N6 atom of AMP and a side-chain oxygen atom of Asp33 and between the N6 atom of AMP and the C1 atom of GSPP are indicated by yellow dashed lines and labeled in Å.

### TvAPT exhibits broad prenyl-acceptor promiscuity

In the substrate-analogue-bound structure of TvAPT_20per, the 5’-phosphate group of AMP is positioned to form hydrogen bonds with Arg34 and the side-chain hydroxyl groups of Ser100 and Ser102, with the latter two residues forming an SXS motif (Figs. 3D and 3G). This motif is conserved among AMP-prenylating enzymes but is absent from their tRNA-prenylating counterparts ^40,41^, suggesting that it may contribute to the preference of TvAPT for nucleoside 5’-monophosphates (Fig. 3D and Supplementary Fig. 10). Guided by these structural observations, we examined adenine-containing substrates with different numbers or arrangements of phosphate groups. Although determinations of kinetic parameters or comparisons of initial rates would have been preferable, reliable kinetic measurements were not feasible for substrates exhibiting low activity. We therefore quantitatively compared their conversion percentages after 16 h under standardized reaction conditions. No prenylated products derived from intact ATP (**5**) or cyclic AMP (**7**) were detected, in contrast to the nearly complete conversion of AMP (**1**), indicating that either the bulky 5’-triphosphate group or 3’,5’-cyclization prevents productive substrate recognition (Fig. 4 and Supplementary Fig. 14). ADP (**6**) was accepted as a substrate, with 35.8% conversion, indicating that a 5’-diphosphate group can be accommodated, although less efficiently than the 5’-monophosphate group of AMP. ATP was extensively dephosphorylated to ADP in a TvAPT-dependent manner, whereas only minor dephosphorylation of ADP to AMP was observed. Similar nucleotide dephosphorylation was previously observed during the characterization of NolPT1, a homologous adenylate isopentenyltransferase that catalyzes the N6-dimethylallylation of AMP using DMAPP^27^ (Supplementary Fig. 14). By contrast, adenosine (**8**), which lacks a 5’-phosphate group, showed only 2.7% conversion. Together, these results support the importance of the 5’-monophosphate group for efficient substrate recognition by TvAPT.

To further examine whether TvAPT could act on adenines within nucleic acids, we evaluated a panel of 6-mer single-stranded oligodeoxynucleotides (AAAAAA, ATTTTT, TTTTTA, and TTTATT, **12-15**). Prenylated products were detected for AAAAAA (**12**; 0.6% conversion) and TTTTTA (**14**; 0.1% conversion), indicating that 3’-terminal adenines can be modified, albeit with very low efficiency (Fig. 4 and Supplementary Fig. 15). Given that the TvAPT genomic neighborhood harbors genes related to the metabolism of NAD (**16**), FAD (**17**), CoA (**18**), GMP, and cyclic di-GMP (**19**), we assessed these cofactors as candidate substrates (Fig. 1C and Supplementary Fig. 3). Assays revealed marginal prenylation activity toward NAD (2.1% conversion) and FAD (3.7% conversion), but no detectable activity for CoA or cyclic di-GMP (Fig. 4 and Supplementary Fig. 14). Thus, while bulky substituents distal to the 5’-diphosphate are partially tolerated, additional modifications, such as the 3’-phosphate of CoA, completely abolish this already marginal activity.

The ribose 2’-hydroxyl group forms a hydrogen bond with the backbone carbonyl of Thr171, whereas no direct interaction with the 3’-hydroxyl group was observed (Fig. 3G). To examine the tolerance of TvAPT toward modifications at the 2’ and 3’ positions, we tested 2’-deoxy-AMP (dAMP, **2**), fludarabine monophosphate (**3**), and 2’,3’-O-trinitrophenyl AMP (TNP-AMP, **4**). Fludarabine monophosphate is an AMP analogue containing a 2-fluoroadenine base and an arabinose sugar with the configuration at C2’ inverted relative to ribose. Although the kinetic analysis showed that dAMP had a higher *K*_□,app_ and a lower *k*_cat,app_ than AMP, both dAMP and fludarabine monophosphate reached complete conversion under the 16-h endpoint assay conditions (Fig. 4 and Supplementary Fig. 6). By contrast, TNP-AMP showed only 2.1% conversion, indicating that TvAPT tolerates the removal or inversion of the 2’-hydroxyl group more readily than a bulky substituent spanning the 2’ and 3’ positions.

Finally, we examined substitutions on the adenine base. GMP (**11**), which lacks the target N6 amino group, and εAMP (**10**), in which the N6 nitrogen is incorporated into an additional five-membered ring, were inactive. TvAPT also converted N6-methyl-AMP (**9**) into both N6-methyl-N6-geranyl-AMP and the demethylated product N6-geranyl-AMP (Supplementary Fig. 12). When only the formation of the methyl-retaining product was considered, the conversion of N6-methyl-AMP to N6-methyl-N6-geranyl-AMP was 18.5%.

**Figure 4.**
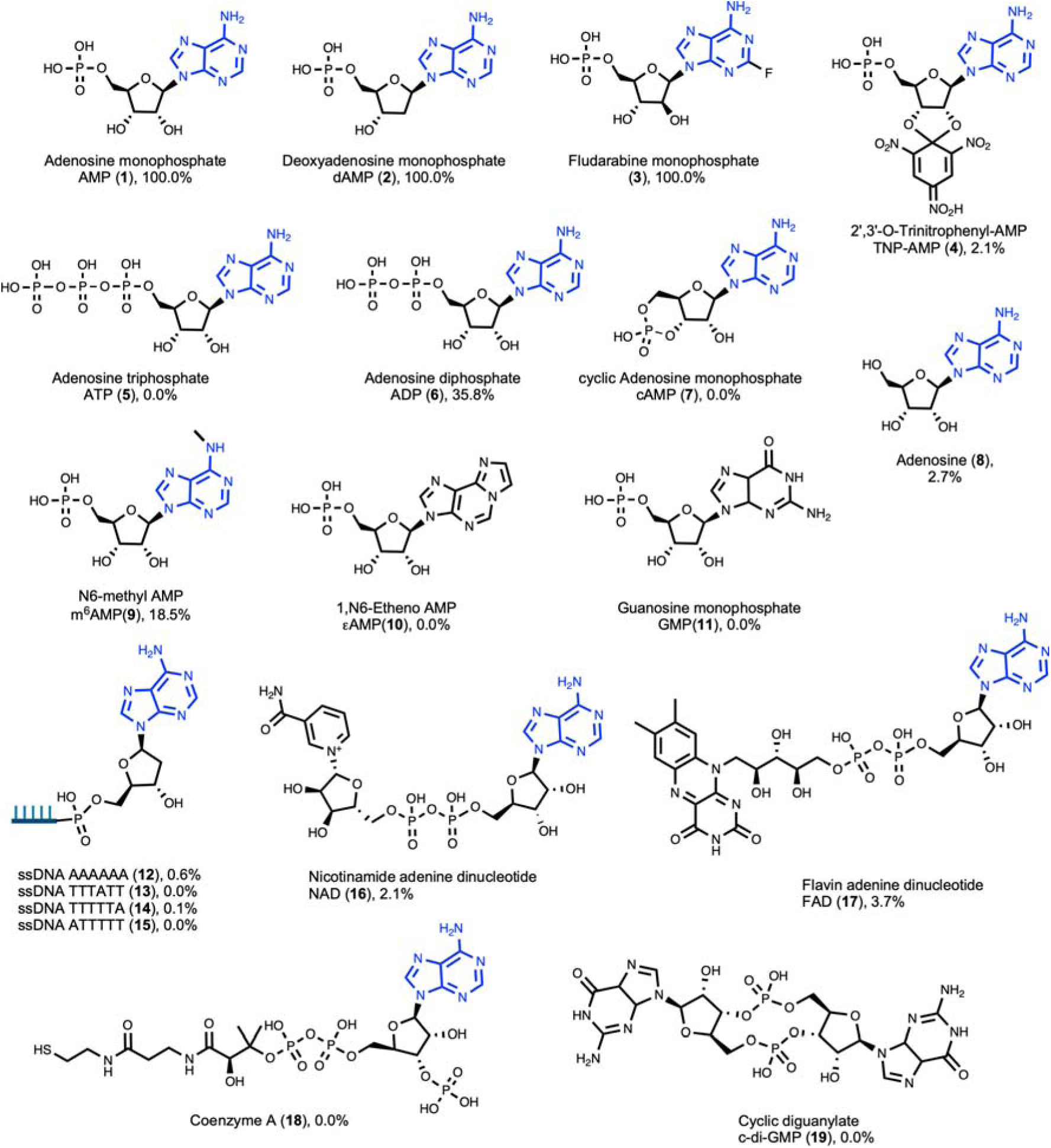
Substrate scope of TvAPT toward adenine-containing substrates. Chemical structures of the evaluated compounds are shown, with the adenine moiety highlighted in blue. The *in vitro* conversion percentages for each substrate are indicated below the corresponding structure.

### Long-chain prenylation enhances the membrane permeability of an AMP analogue

Although nucleotide-based compounds are attractive therapeutic candidates, they typically exhibit poor membrane permeability owing to their negatively charged phosphate moiety and limited overall hydrophobicity^44^. We hypothesized that enzymatic prenylation could improve cellular entry, as observed for engineered peptides, in which the C-terminal installation of a C15 farnesyl group promotes the diffusive cellular uptake of otherwise impermeable species^45^. To test this, we enzymatically synthesized TNP-AMP derivatives bearing C5-, C10-, or C15-prenyl chains and evaluated their cell permeability (Fig. 5A and Supplementary Fig. 16). To obtain sufficient quantities for these cellular assays, we increased the conversion yield from marginal levels to over 50% by increasing the enzyme concentration more than threefold and extending the reaction time from overnight to 3 days. Following a 1.5-h incubation of the HPLC-purified compounds with mouse embryonic fibroblasts (MEFs) at 37°C, intracellular fluorescence was analyzed by microscopy. Cellular uptake was quantified as the fraction of the bright-field-defined cell area exhibiting positive fluorescence. Unmodified TNP-AMP displayed a negligible fluorescence-positive area (0.5%; Figs. 5B and C), confirming its poor membrane permeability. In contrast, the C10- and C15-prenylated derivatives exhibited substantially higher intracellular fluorescence (15.6% and 12.2%, respectively), whereas the C5-prenylated analogue showed only a modest increase (4.6%; Figs. 5B and C).

Lipid-modified deoxyribonucleoside triphosphates (dNTPs), featuring nucleobases chemically conjugated to monoacyl, diacyl, or cholesterol moieties, have been previously reported^21^. Upon PCR-mediated incorporation into DNA strands, these modified nucleotides enable the resulting nucleic acids to function as stable membrane anchors. Comparative studies indicated that the membrane anchoring is strongly influenced by the lipid architecture, with diacyl, unsaturated acyl, and cholesterol modifications generally proving more effective^46^. However, prenyl groups, which feature repeating double bonds and branched methyl groups, have not been systematically evaluated in this context. Our findings suggest that prenylated TNP-AMP preferentially undergoes cellular internalization, rather than stable membrane anchoring. This likely reflects both the significantly smaller size of TNP-AMP relative to nucleic acids, which lowers the free-energy barrier for membrane translocation, and the distinctive membrane-partitioning properties of prenyl groups^47^. Consistent with this model, prenylation promoted the partitioning of molecules into more loosely packed, non-raft membrane regions that are generally more permissive to translocation than the rigid domains favored by cholesterolylation or acylation^48^.

**Figure 5.**
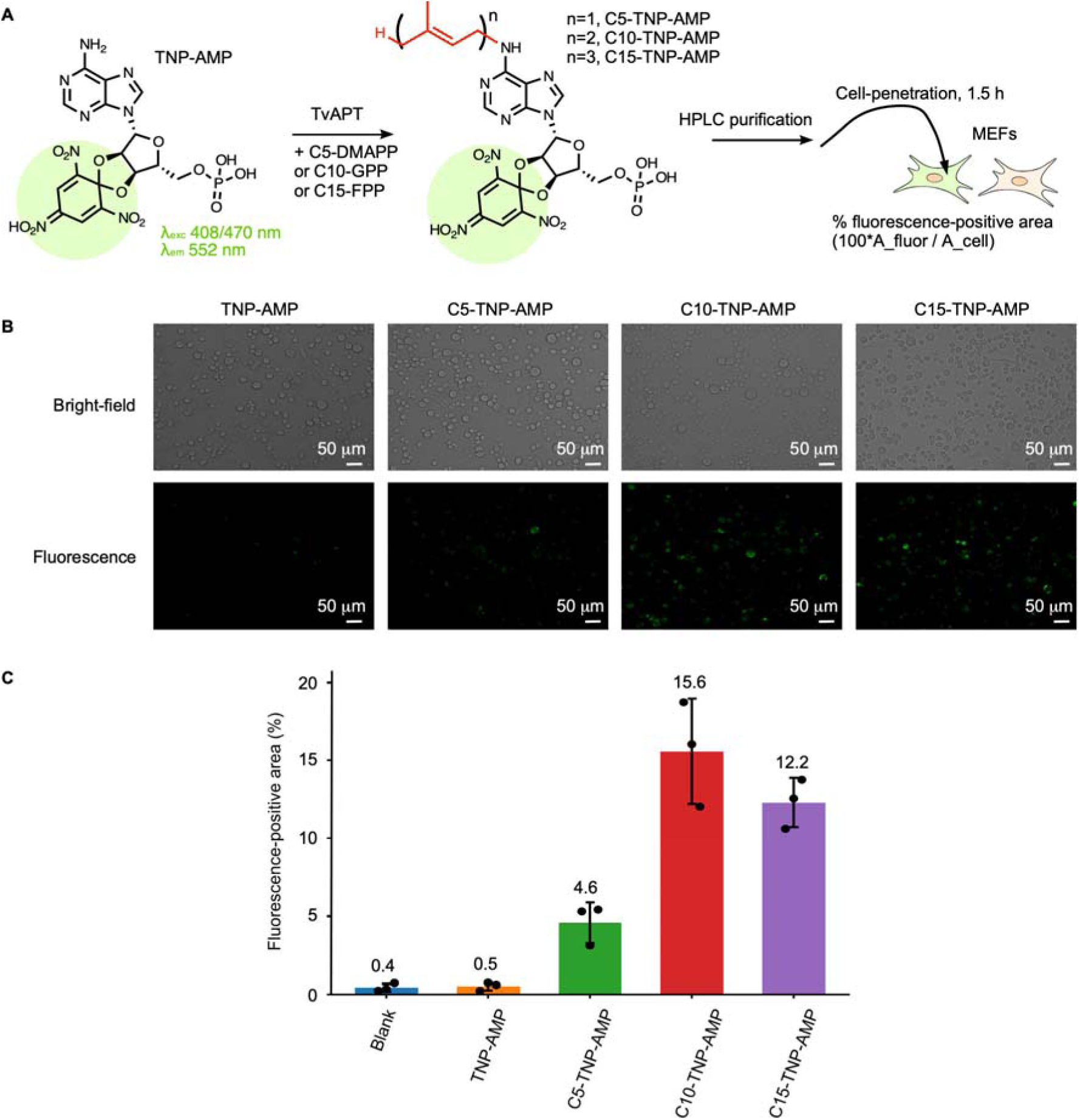
Long-chain prenylation enhances the cellular uptake of TNP-AMP. (A) Schematic representation of the cell permeability assay. The fluorescent TNP moiety is highlighted in green. (B) Fluorescence microscopy of MEF cells incubated with TNP-AMP derivatives bearing C5-, C10-, and C15-prenyl chains. Bright-field (upper panels) and corresponding fluorescence images (lower panels) are shown. (C) Quantification of the fluorescence-positive areas. Bars represent the mean percentage of fluorescence-positive area ± s.d. from three independent experiments. Mean values are shown above the bars.

### Long-chain cytokinin analogues suppress root hair formation in *Arabidopsis thaliana*

Cytokinins, including C5-iP and related N6-derivatives, are master regulators of plant growth and development, orchestrating processes such as leaf senescence, shoot branching, root development, and stress responses ^49^. These phytohormones bind to histidine kinase (HK) receptors at the plasma membrane and endoplasmic reticulum, regulating downstream transcription factors via a phosphorelay signaling cascade (Fig. 6E) ^50,51^. To evaluate the biological effects of extended prenyl chains, we enzymatically synthesized long-chain cytokinin analogues. Specifically, AMP was prenylated by TvAPT, and subsequent treatment with LOG from *Pseudomonas aeruginosa* PAO1 (PaLOG)^52^ yielded the targeted analogues C5-iP, C10-gP, and C15-fP (Fig. 6A and Supplementary Fig. 17). Four-day-old *Arabidopsis thaliana* seedlings were treated with these analogues, using 6-benzylaminopurine (6-BP) as a positive control, and root hair formation was assessed 24 h later. Root hair density was calculated as the number of root hairs per millimeter. Root hairs visible on one side of the primary root were counted within a 0.5-mm segment extending shootward from the first visible root hair bulge near the root tip. As expected, 6-BP and C5-iP markedly increased root hair density from 24.1 root hairs mm ¹ in the untreated control to 83.5 and 65.4 root hairs mm ¹, respectively (Fig. 6B and C). By contrast, the long-chain derivatives C10-gP and C15-fP did not promote root hair formation, with densities of 15.8 and 18.7 root hairs mm ¹, respectively, which were both lower than that of the untreated control.

Cytokinin signaling is modulated by type-B ARRs, which drive responsive transcription, and type-A ARRs (e.g., *ARR5* and *ARR15*), which exert negative feedback control (Fig. 6E)^53^. Because *ARR15* overexpression is known to reduce the sensitivity to exogenous cytokinins^54^, we quantified *ARR5* and *ARR15* transcript levels at 2 hours and 2 days post-treatment to investigate the mechanism underlying this root hair suppression. Notably, robust early induction of both genes was observed only with C15-fP at the 2-hour time point, consistent with its modest inhibitory effect on root hair formation (Fig. 6D). In contrast, no significant upregulation was detected for either C10-gP or C15-fP at 2 days under the tested conditions.

The histidine kinase receptor AHK4 has been reported to mediate *ARR15* induction^54^. To gain insight into whether the potent upregulation of type-A ARR expression by C15-fP reflects stronger binding to HK receptors, we performed *in silico* binding analyses of AHK4 with several cytokinin derivatives (Supplementary Table 1 and Supplementary Fig. 18). Although a direct comparison of the experimentally determined binding constants is complicated by differences in methodology, C5-iP reportedly binds AHK4 more strongly than 6-BP^55,56^. In our structural predictions, the estimated binding scores followed the order C10-gP > C5-iP > C15-fP > 6-BP. Inspection of the AHK4 binding site suggested that the accommodation of C15-fP would require kinking of the farnesyl chain (Supplementary Fig. 18). Moreover, the bulky farnesyl group appeared to displace the adenine moiety, further implying suboptimal interactions between C15-fP and AHK4. Taken together, these observations suggest that the effect of C15-fP on *ARR15* expression may be related to altered distribution within plant cells or tissues, rather than enhanced binding to AHK4^49^.

**Figure 6.**
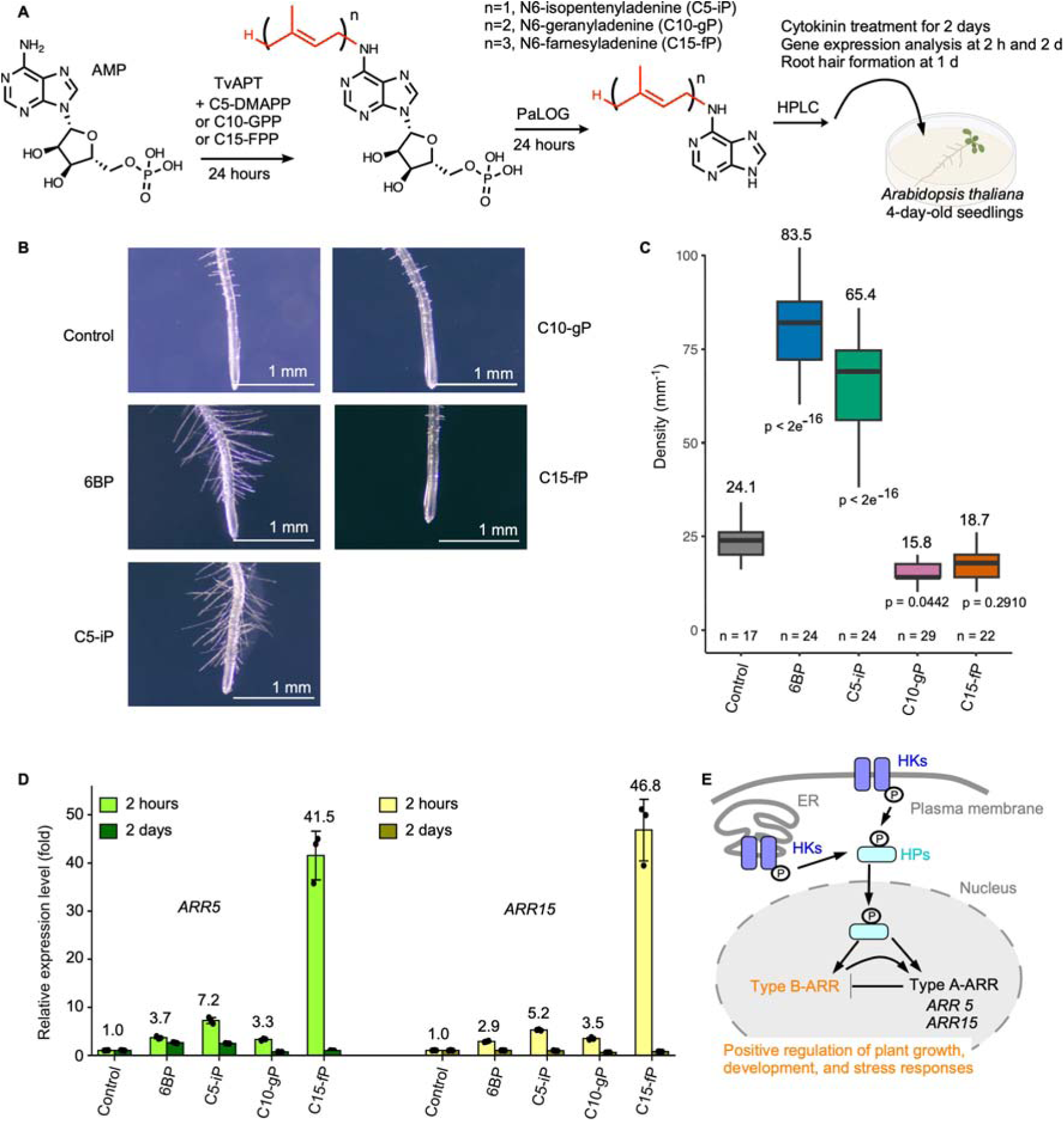
Biological evaluation of long-chain cytokinin analogues in *Arabidopsis thaliana*. (A) Schematic illustration of the chemoenzymatic synthesis of N6-prenyladenine analogues and their subsequent biological evaluation. AMP was prenylated by TvAPT using DMAPP, GPP, or FPP and then dephosphorylated by PaLOG. Four-day-old *A. thaliana* seedlings were treated with the resulting C5-iP, C10-gP, or C15-fP, and gene expression and root hair formation were evaluated at the indicated time points. (B) Representative root hair phenotypes 24 h after mock treatment (control) or treatment with 6-benzylaminopurine (6-BP), N6-isopentenyladenine (C5-iP), N6-geranyladenine (C10-gP), or N6-farnesyladenine (C15-fP). Images are representative of three independent experiments. Scale bars, 1 mm. (C) Quantification of root hair density. Root hairs visible on one side of the primary root were counted within a 0.5-mm segment extending shootward from the first visible root hair bulge and expressed as the number of root hairs per millimeter. Boxes indicate the interquartile range, center lines indicate the median, and whiskers extend to 1.5 times the interquartile range. Mean values are shown above the boxes, and n values shown below the boxes indicate the number of 0.5-mm root segments analyzed. Adjusted *P* values for comparisons with the control were calculated using one-way ANOVA followed by Dunnett’s multiple-comparisons test and are indicated in the graph. (D) Relative transcript levels of *ARR5* and *ARR15* at 2 h and 2 d after treatment, as quantified by RT-qPCR. Data are presented as the mean fold change ± s.d. from three independent experiments. Mean values at 2 h are shown above the bars. (E) Simplified schematic of the cytokinin signaling cascade. HKs, histidine kinases; HPs, histidine phosphotransfer proteins. Nuclear translocation of phosphorylated HPs regulates type-A and type-B ARR transcriptional regulators, with type-A ARRs mediating a negative-feedback loop.

## Discussion

In this study, we identified TvAPT as an adenine prenyltransferase capable of transferring extended prenyl chains to the N6 position of adenine, a transformation inaccessible to canonical C5-dimethylallyltransferases. Structural analyses revealed that this expanded donor scope is dictated by an enlarged prenyl-binding pocket, and mutational studies demonstrated that the prenyl-donor preference can be tuned via a single-residue substitution. Beyond AMP, TvAPT accommodated several adenine-containing substrates, with a clear preference for nucleoside 5’-monophosphates. However, preparative synthesis, particularly with less-preferred substrates, currently requires relatively high enzyme loading and prolonged reaction times. Furthermore, soluble production of wild-type TvAPT requires chaperone co-expression (Supplementary Fig. 20). Future protein-engineering efforts aimed at improving catalytic efficiency and soluble protein production will therefore be important for broader synthetic applications.

The weak but detectable activity of TvAPT toward diverse adenine-containing acceptors suggests that its acceptor scope could be further expanded by directed evolution. TvAPT also catalyzed the site-selective prenylation of the 3’-terminal adenine in short single-stranded oligodeoxynucleotides. This provides, to our knowledge, the first example of the enzymatic prenylation of short ssDNA oligonucleotides. Notably, adenosine, which bears a free 5’-hydroxyl group, was a relatively poor substrate, whereas the conversion increased markedly when a 5’-monophosphate was present. A similar principle may apply to single-stranded oligodeoxynucleotides, where capping the 5’-hydroxyl group with a phosphate could enhance substrate recognition and catalytic efficiency. The inability of TvAPT to modify internal nucleobases likely reflects the absence of the dedicated nucleic acid-binding domains found in canonical tRNA prenyltransferases (Supplementary Fig. 10). Therefore, the introduction of long-chain prenyl groups at internal positions may require engineering canonical tRNA prenyltransferases so that they can accept extended prenyl donors while retaining their nucleic acid-recognition properties. Given the growing interest in membrane-localized RNA^57,58^ and DNA ^59^ devices, such engineered biocatalysts could facilitate the development of novel oligonucleotides with tunable membrane behaviors.

Biologically, our results raise the possibility that TvAPT participates in the biosynthesis of previously unrecognized cytokinin-like signaling molecules in *Trichormus variabilis* NIES-23. The *T. variabilis* genome encodes both a LOG-family protein and a CKX-like protein, consistent with active cytokinin metabolism, yet lacks recognizable HK or HP receptor genes (Supplementary Fig. 19). Although the physiological roles of these metabolites remain unknown, long-chain adenine derivatives may influence interspecies interactions between cyanobacteria and plants ^60^. More broadly, our findings support the emerging paradigm that side-chain diversification fundamentally redirects cytokinin activity ^61^. Consistent with this view, the long-chain TvAPT-derived analogues failed to promote root hair formation in *A. thaliana*. A related mechanism has been reported in the plant pathogen *Rhodococcus fascians*, where the AIPT enzyme catalyzes the N6-diethylallylation of AMP to produce a cytokinin-like root growth inhibitor ^60^. Notably, the critical residue at position 142 is a bulky methionine in *R. fascians* AIPT, whereas cyanobacterial APTs invariably feature an alanine at this equivalent position (Supplementary Fig. 19). This structural divergence suggests that different prenyltransferase lineages have independently evolved alternative structural solutions to accommodate expanded side-chain diversity.

## Methods

### Bioinformatic analysis

Sequence similarity networks were generated using the online Enzyme Function Initiative-Enzyme Similarity Tool (EFI–EST)^26^. Adenylate dimethylallyltransferases from the InterPro families IPR002648 and IPR039657 were used as queries. Nodes were clustered using an alignment score threshold of 60, which corresponds to approximately 35% sequence identity. The resulting network was visualized using Cytoscape version 3.8.0^62^. To analyze genomic contexts, the XGMML output file from EFI-EST was processed using EFI-GNT, and the resulting genome neighborhood networks were examined manually. For phylogenetic analysis, sequences of representative members from the adenylate dimethylallyltransferase family and the TvAPT-containing clade were aligned using MAFFT. A phylogenetic tree was subsequently constructed from this alignment.

### Plasmid construction

Using the pET-28b(+) vector, the sequences encoding TvAPT and tzs were inserted between the NheI and XhoI sites, TvAPT_20per between the NdeI and XhoI sites, and PaLOG between the NcoI and XhoI sites to express the proteins with N-terminal His tags, except for PaLOG, which was expressed with a C-terminal His tag. All genes were codon-optimized for expression in *E. coli* and synthesized by GenScript Japan. Site-directed mutagenesis to generate the TvAPT A142M and tzs M142A variants was performed using a KOD Mutagenesis Kit (Toyobo) with the primers listed in Supplementary Table 2. Sequence fidelity for all constructs was verified by DNA sequencing using standard T7 promoter and T7 terminator primers. The corresponding amino acid sequences for TvAPT, TvAPT_20per, tzs, and PaLOG are provided in Supplementary Table 3.

### Protein expression and purification

Recombinant proteins were expressed in *E. coli* BL21(DE3), grown at 37°C in LB medium. For TvAPT, tzs, and their variants, cells were co-transformed with the pG-KJE8 chaperone plasmid (Takara) and cultured with 30 µg/mL kanamycin, 40 µg/mL chloramphenicol, 5 ng/mL tetracycline, and 0.12% arabinose. For PaLOG, cells were cultured with 30 µg/mL kanamycin alone. At an OD_600_ of 0.6, expression was induced with 0.5 mM IPTG, followed by overnight incubation at 20°C. Cells were harvested by centrifugation (5,000 × g), resuspended in buffer A (50 mM Tris-HCl, pH 7.5, 150 mM NaCl, and 15% glycerol), and lysed by sonication. After centrifugation (9,000 × g, 10 min), the clarified lysates were loaded onto HisPur Ni-NTA resin (Thermo Scientific) pre-equilibrated with buffer A.

The resin was washed with buffer A containing 20 mM imidazole. For PaLOG, the protein was then eluted with 400 mM imidazole in buffer A and used directly for enzymatic assays. For TvAPT and tzs variants, an additional chaperone-removal wash (100 mM Tris-HCl, pH 7.5, 100 mM KCl, 20 mM MgCl_2_, 25% glycerol, 500 mM sucrose, and 5 mM ATP) was performed prior to elution with 400 mM imidazole. The eluted TvAPT and tzs fractions were adjusted to 300 mM Na_2_SO_4_, analyzed by SDS-PAGE (Supplementary Fig. 16), and further purified by size-exclusion chromatography on a Superdex 200 Increase column (Cytiva) equilibrated with buffer B (20 mM HEPES-NaOH, pH 7.5, 300 mM Na_2_SO_4_, and 10% glycerol). Protein concentrations were determined spectrophotometrically at 280 nm. For metal-dependence assays, purified TvAPT was treated with 50 mM EDTA for 4 h and dialyzed against buffer B.

### Structural determination of TvAPT_20per

The amino acid sequence of TvAPT was analyzed using ESM-scan^36^. For each position, ESM scores for all 20 possible amino acid substitutions were calculated and averaged. The 25 positions exhibiting the highest mean scores, representing approximately 10% of the 244-residue sequence, were selected for mutation. These targeted sites were replaced with residues proposed by LigandMPNN^37^, yielding 20 redesigned sequence variants. These variants were then integrated into a single consensus sequence via ancestral reconstruction using MAFFT2ASR (https://github.com/shognakano/MAFFT2ASR). The resulting engineered construct, designated TvAPT_20per, was expressed and purified as described above.

TvAPT_20per was crystallized in the product-bound and substrate-analogue-bound forms by sitting-drop vapor diffusion at 23°C. For the product-bound form, equal volumes of protein solution (12.4 mg/mL TvAPT_20per in 20 mM HEPES–NaOH, pH 7.5, 300 mM Na SO , 5 mM MgCl , 1 mM GPP, and 10% glycerol) and reservoir solution (1.9 M ammonium sulfate and 100 mM MES–NaOH, pH 6.5) were mixed. For the substrate-analogue-bound form, equal volumes of protein solution (14.2 mg/mL TvAPT_20per in 20 mM HEPES–NaOH, pH 7.5, 300 mM Na SO , 5 mM MgCl , 2 mM GSPP, 2 mM AMP, and 10% glycerol) and reservoir solution (1.0 M sodium citrate, pH 6.2) were mixed. Crystals were briefly soaked in the corresponding reservoir solution supplemented with 25% (v/v) glycerol and flash-cooled in liquid nitrogen before X-ray data collection. Diffraction data were collected at beamline BL-5A of the Photon Factory (Tsukuba, Japan) and processed using XDS^63^. Initial phases were determined by molecular replacement in MOLREP ^64,65^, utilizing the predicted structural model (described below) as the search template. Iterative cycles of manual model building and crystallographic refinement were performed using Coot^65^ and REFMAC^66^, respectively. Data collection and refinement statistics are summarized in Supplementary Table 4.

### Structure predictions

Structural models of TvAPT, tzs, and AHK4 (residues 149-418) were predicted using Boltz-2^34^. The amino acid sequences were provided as inputs, together with the SMILES strings of FPP for TvAPT, DMAPP for tzs, and the cytokinin derivatives for AHK4. The SMILES strings for AMP and Mg^2+^ were also included for TvAPT and tzs. For the TvAPT_20per model used in molecular replacement, only the amino acid sequence was supplied. All predictions were performed using default parameters. The resulting predicted local distance difference test (pLDDT) confidence scores are shown in Supplementary Fig. 5.

### Prenylation assay

Reactions were typically performed in a 50 µL volume containing 30 µM enzyme (TvAPT, tzs, or their variants), 400 µM prenyl diphosphate, 100 µM prenyl acceptor, and 1 mM MgCl_2_ in buffer A (50 mM Tris-HCl, pH 7.5, 150 mM NaCl, and 15% glycerol). For prenyl-donor competition assays, the reaction mixture contained an equimolar mixture of DMAPP, GPP, FPP, and GGPP, each at a final concentration of 100 µM. For acceptor-scope evaluations, the following substrates were tested: AMP (adenosine 5’-monophosphate), ATP (adenosine 5’-triphosphate), ADP (adenosine 5’-diphosphate), adenosine, dAMP (2’-deoxyadenosine 5’-monophosphate), cAMP (3’,5’-cyclic adenosine monophosphate), εAMP (1,N6-ethenoadenosine 5’-monophosphate), N6-methyl-AMP (N6-methyladenosine 5’-monophosphate), fludarabine monophosphate, GMP (guanosine 5’-monophosphate), TNP-AMP (2’,3’-O-(2,4,6-trinitrophenyl)adenosine 5’-monophosphate), NAD (nicotinamide adenine dinucleotide), FAD (flavin adenine dinucleotide), coenzyme A, cyclic di-GMP, and 6-mer ssDNAs (AAAAAA, ATTTTT, TTTTTA, or TTTATT). For metal-dependence assays, MgCl_2_ was substituted with 1 mM ZnCl_2_, NiCl_2_, MnCl_2_, CuCl_2_, CaCl_2_, or CoCl_2_.

Following a 16-h incubation at room temperature, the reaction mixtures were analyzed by HPLC. Chromatography was performed on a COSMOSIL 5C18-AR-II column (4.6 mm × 100 mm; Nacalai Tesque) at a flow rate of 1 mL/min. The mobile phase comprised solvent A (0.1 M triethylammonium acetate, pH 7.0) and solvent B (acetonitrile), utilizing a linear gradient from 0% to 100% solvent B over 20 min. Analytes were detected at 260 nm. Eluted peaks of interest were collected and analyzed via MALDI-TOF MS (Bruker). For mass spectrometry sample preparation, 1 µL of the collected fraction was spotted onto a target plate and dried. This was followed by the sequential addition of 0.5 µL of 2,4,6-trihydroxyacetophenone (70 mg/mL in 60% acetonitrile) and 0.5 µL of ammonium citrate (50 mg/mL in 50% acetonitrile). After air-drying, mass spectra were acquired in positive-ion mode.

### Steady-state kinetic assays

Reactions were typically performed in a 50 µL volume, containing 2 μM enzyme for wild-type TvAPT with dAMP or TvAPT_20per with AMP, or 0.4 μM enzyme for wild-type TvAPT with AMP, 500 µM prenyl diphosphate, variable concentrations of AMP or dAMP, and 1 mM MgCl in buffer A (50 mM Tris-HCl, pH 7.5, 150 mM NaCl, 300 mM Na SO , and 15% glycerol). A 30-min reaction time was selected because product formation was approximately linear over this interval. After incubation for 30 min, the reactions were quenched by adding an equal volume of 0.1 M EDTA solution and incubating the samples on ice. The quenched samples were analyzed by HPLC as described above. All assays were performed in triplicate. Product formation was quantified by HPLC peak integration, and apparent product concentrations were estimated from the product fraction of the combined substrate and product peak areas, using the known initial substrate concentration. Apparent reaction velocities were calculated from product accumulation after 30 min, and individual triplicate data points were fitted to the Michaelis–Menten model by nonlinear least-squares regression. Because the assay was performed at a fixed GPP concentration of 500 µM, corresponding to a 250- to 1,000-fold molar excess over the enzyme concentration, and was based on a 30-min endpoint assay, the resulting kinetic parameters are reported as apparent values.

### NMR spectroscopy

A 5 mL reaction mixture, containing 30 µM TvAPT, 200 µM C10-GPP, 1 mM MgCl_2_, and 1 mM AMP in buffer A, was incubated overnight at room temperature. Preparative HPLC purification was performed under the same conditions as described above, except that a COSMOSIL 5C18-AR-II column (10 mm × 250 mm, Nacalai Tesque) was used at a flow rate of 2.5 mL/min and elution was carried out with a linear gradient from 0% to 100% solvent B over 40 min. The fraction containing prenylated AMP was collected and concentrated to dryness using a centrifugal concentrator. The NMR sample (0.25 mL) contained 0.6 mM prenylated AMP dissolved in 100% D_2_O. NMR spectra were recorded at 298 K on a Bruker Avance 500 MHz spectrometer equipped with a TXI cryoprobe. ^1^H and ^13^C resonance assignments were obtained from ^1^H-^13^C gHSQC and ^1^H-^13^C HMBC spectra. A one-dimensional ^1^H NMR spectrum was acquired using a standard presaturation pulse program for solvent suppression. NMR data were processed and analyzed using the POKY suite^67^. The ^1^H and ^13^C resonance assignments are provided in Supplementary Table 5.

### High-resolution mass spectrometry

N6-Geranylated AMP, prepared in 100% D_2_O for NMR measurements, was diluted with 300 mM ammonium acetate (pH 6.8) to a final concentration of 500 μM. Approximately 2–5 μL of the sample was loaded into a custom-made gold-coated glass capillary and analyzed by nanoflow electrospray ionization mass spectrometry, using a SYNAPT G2-Si HDMS mass spectrometer (Waters) operated in the negative-ion mode. Measurements were performed under the following instrument conditions: capillary voltage, 1.13 kV; sampling cone voltage, 150 V; source offset, 150 V; trap gas flow rate, 5 mL/min; trap collision energy, 4 V; and transfer collision energy, 2 V. The spectra were calibrated using a 2 mg/mL cesium iodide solution in 50% (v/v) aqueous 2-propanol and analyzed using the MassLynx software (Waters).

### Enzymatic synthesis of prenylated TNP-AMP and cell permeability assay

Preparative-scale reactions (0.7 mL), containing 90 µM TvAPT, 100 µM TNP-AMP, 400 µM prenyl diphosphate (C5-DMAPP, C10-GPP, or C15-FPP), and 1 mM MgCl_2_ in buffer A, were incubated for three days at room temperature. Preparative HPLC purification was performed using the conditions described above. Fractions containing the target prenylated TNP-AMP derivatives were pooled, evaporated to dryness via vacuum centrifugation, and dissolved in water to a final concentration of 400 µM.

Mouse embryonic fibroblasts (MEFs) were cultured in 96-well plates in Dulbecco’s modified Eagle medium (DMEM; high glucose) containing L-glutamine and phenol red, supplemented with 10% fetal bovine serum (FBS; Biowest), 100 U/mL penicillin G, 100 µg/mL streptomycin, and 0.25 µg/mL amphotericin B. Prior to treatment, the cells were washed with Hanks’ Balanced Salt Solution (HBSS). Unmodified TNP-AMP and the purified C5-, C10-, and C15-prenylated derivatives were then added to a final concentration of 2 µM. Mock-treated cells served as negative controls, and all conditions were evaluated in triplicate. Following a 1.5-h incubation, cells were examined using a BZ-X800 all-in-one fluorescence microscope (Keyence). Bright-field and corresponding GFP-channel fluorescence images were acquired at 40× magnification from three independent fields per sample. Fluorescence exposure times were calibrated to ensure no background signal was detected in the negative controls. Image analysis was conducted using the Hybrid Cell Count module of the BZ-X800 Analyzer software. For bright-field images, thresholds were manually adjusted to encompass all cells within the field, and the total cell area (μm^2^) was calculated with the hole-filling function set to “strong.” For fluorescence images, a uniform intensity threshold of 10 was applied to delineate the fluorescence-positive area. Cellular permeability was ultimately quantified as the fraction of the total bright-field cell area exhibiting positive fluorescence.

### Root hair formation assay in *Arabidopsis thaliana*

Preparative-scale reactions (1.5 mL), containing 60 µM TvAPT, 300 µM AMP, 400 µM prenyl diphosphate (C5-DMAPP, C10-GPP, or C15-FPP), and 1 mM MgCl_2_ in buffer A, were incubated for 24 h at room temperature. PaLOG was subsequently added to a final concentration of 60 µM, followed by an additional 24-h incubation at room temperature. The reaction mixtures were analyzed and purified by preparative HPLC, using the conditions described above. Fractions containing the target prenylated adenines were pooled, evaporated to dryness via vacuum centrifugation, and dissolved in dimethyl sulfoxide (DMSO) to a final concentration of 500 µM.

*Arabidopsis thaliana* seeds were germinated and grown on Murashige and Skoog (MS) medium for 4 days before transfer to fresh medium supplemented with 50 nM of each test compound. Treatments included a mock-treated (vehicle) control, 6-benzylaminopurine (6-BP) as a positive control, and the enzymatically synthesized cytokinin analogues C5-iP, C10-gP, and C15-fP. Root hair phenotypes were evaluated by microscopy after 24 h of treatment. Root hair density was calculated as the number of root hairs per millimeter. Root hairs were counted within a 0.5-mm segment of the primary root extending shootward from the first visible root hair bulge near the root tip. Because the seedlings remained on the agar medium during observation, only the root hairs visible on one side of the primary root were counted.

The resulting counts were converted to the number of root hairs per millimeter. Root hair analyses were performed in three independent biological experiments. Statistical comparisons between each treatment group and the control were performed using one-way ANOVA followed by Dunnett’s multiple-comparisons test. For gene expression profiling, whole seedlings were harvested at 2 h or 2 days post-treatment. Total RNA was extracted and subjected to RT-qPCR to quantify the transcript levels of the cytokinin-responsive genes *ARR5* and *ARR15*. All RT-qPCR experiments were performed in biological triplicate.

## Data availability

The data supporting the findings of this study are available from the corresponding authors upon reasonable request. The atomic coordinates and structure factors have been deposited in the Protein Data Bank under accession codes 24WM and 27DP.

## Acknowledgements

This work was supported by JSPS KAKENHI (grant numbers 23K13887 to D.F., 25K22481 to K.T., and 23K04510 to S.I.); the Uehara Memorial Foundation, Morinomiyako Medical Research Foundation, Mishima Kaiun Memorial Foundation, Yakult Bio-Science Foundation, Takano Life Science Research Foundation, and the Koyanagi Foundation (to D.F.); Joint Research of ExCELLS (24EXC350, 25EXC347 and 26EXC308); the NMR Platform supported by MEXT, Japan; and MEXT through the Promotion of Development of a Joint Usage/Research System Project, Spin-L (JPMXP1323015488 and spin24XN012). We thank Drs. Kenichiro Todoroki, Kenji Kojima, and Aogu Furusho (University of Shizuoka) for their technical advice on MALDI-TOF MS analyses. We also thank Dr. Susumu Uchiyama (Osaka University) for his technical advice on high-resolution mass spectrometry analyses. We thank the staff of the Photon Factory for X-ray data collection (proposal number 2024G03).

## Competing interests

The authors declare no competing interests.

## Notes

### Competing Interest Statement

The authors have declared no competing interest.

### Summary of Updates

This version has been revised to include additional biochemical, structural, and mass spectrometric analyses. New data include a substrate analogue bound structure of TvAPT 20per, expanded kinetic and metal dependence analyses, additional substrate scope experiments, and high resolution mass spectrometry of the enzymatic product. The figures, supplementary information, and discussion have been updated accordingly.

