## Supplementary Information for "A cyanobacterial adenine prenyltransferase enables longer-chain N6 prenylation"

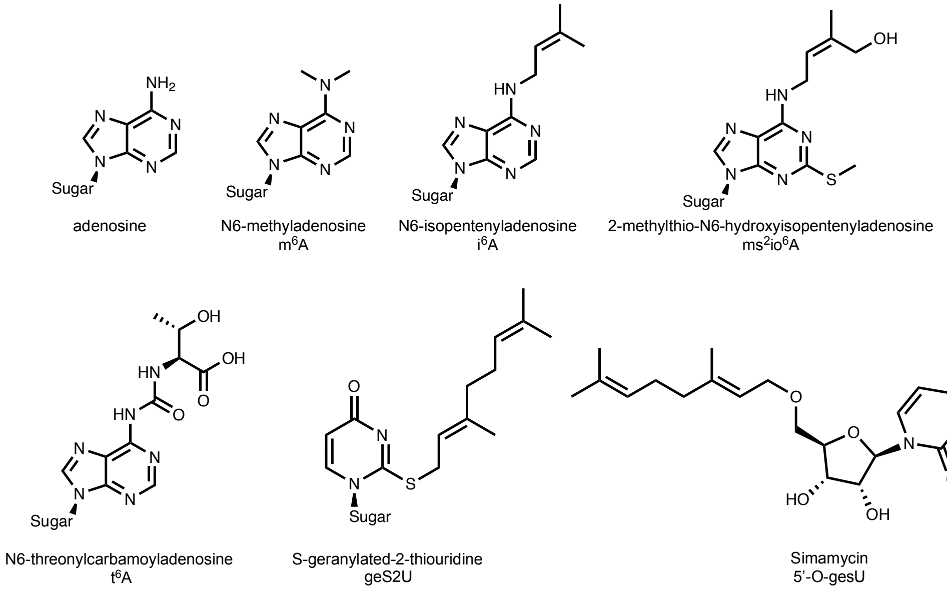


**Supplementary Figure 1.** Chemical structures of N6-modified adenines, along with geranylated uridine nucleosides.

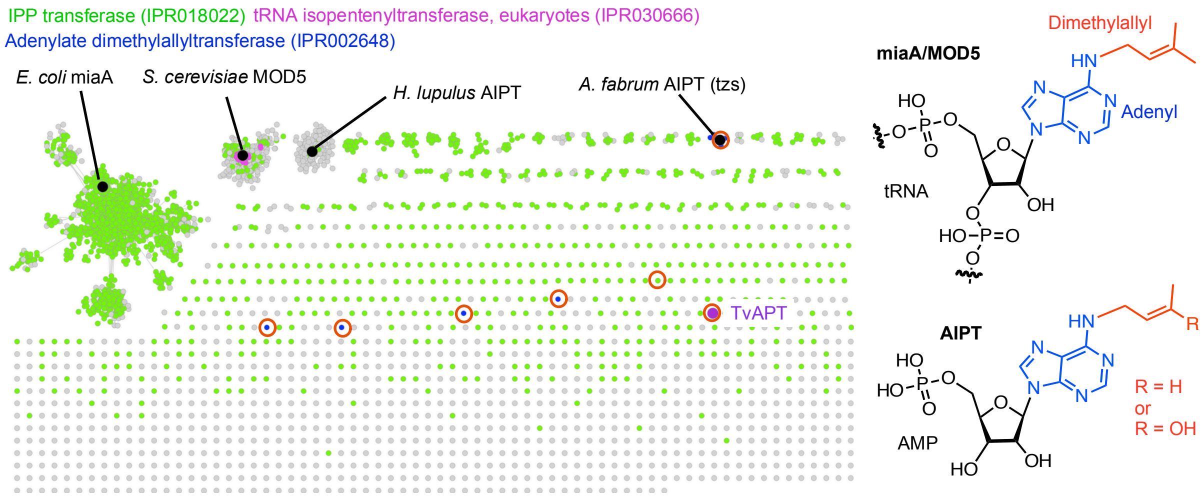


**Supplementary Figure 2.** Sequence similarity network of adenine prenyltransferase homologues. The network was generated using EFI-EST. Clusters corresponding to bacterial tRNA isopentenyltransferases (IPR018022), eukaryotic tRNA isopentenyltransferases (IPR030666), and adenylate dimethylallyltransferases (IPR002648) are indicated in green, purple, and blue, respectively. TvAPT is highlighted in purple. The chemical structures on the right show representative products associated with the indicated sequence clusters. The cluster circled in orange contains bacterial adenylate prenyltransferases closely related to TvAPT and was subjected to the phylogenetic analysis shown in Supplementary Fig. 19.

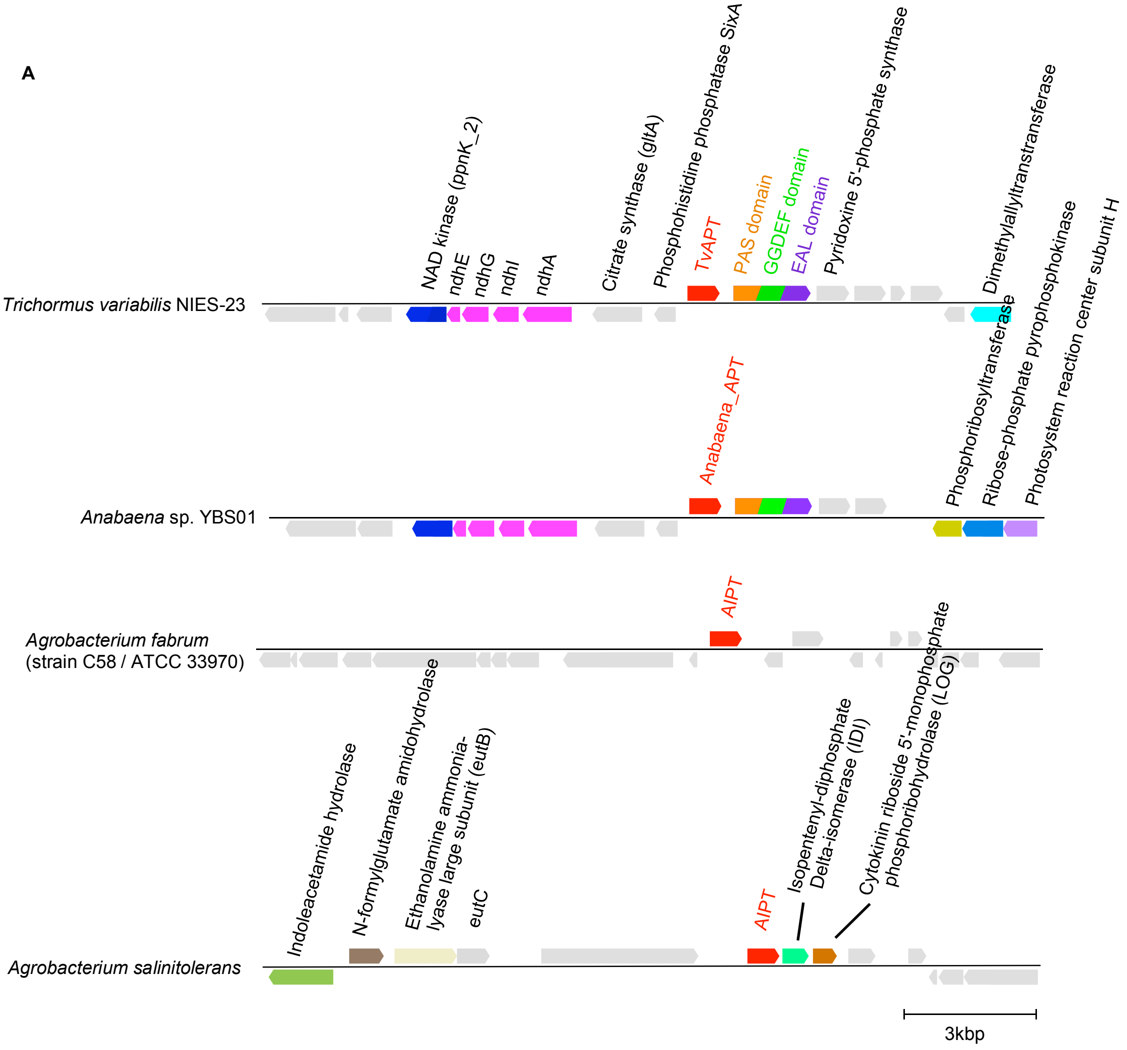
A
Trichormus variabilis NIES-23
NAD kinase (ppnK_2)
ndhE
ndhG
ndhI
ndhA
Citrate synthase (gltA)
Phosphohistidine phosphatase SixA
TvAPT
PAS domain
GGDEF domain
EAL domain
Pyridoxine 5'-phosphate synthase
Dimethylallyltranstransferase
Anabaena sp. YBS01
Anabaena_APT
Phosphoribosyltransferase
Ribose-phosphate pyrophosphokinase
Photosystem reaction center subunit H
Agrobacterium fabrum
(strain C58 / ATCC 33970)
AIPT
Agrobacterium salinitolerans
Indoleacetamide hydrolase
N-formylglutamate amidohydrolase
Ethanolamine ammonia-lyase large subunit (eutB)
eutC
AIPT
Isopentenyl-diphosphate Delta-isomerase (IDI)
Cytokinin riboside 5'-monophosphate phosphoribohydrolase (LOG)
3kbp

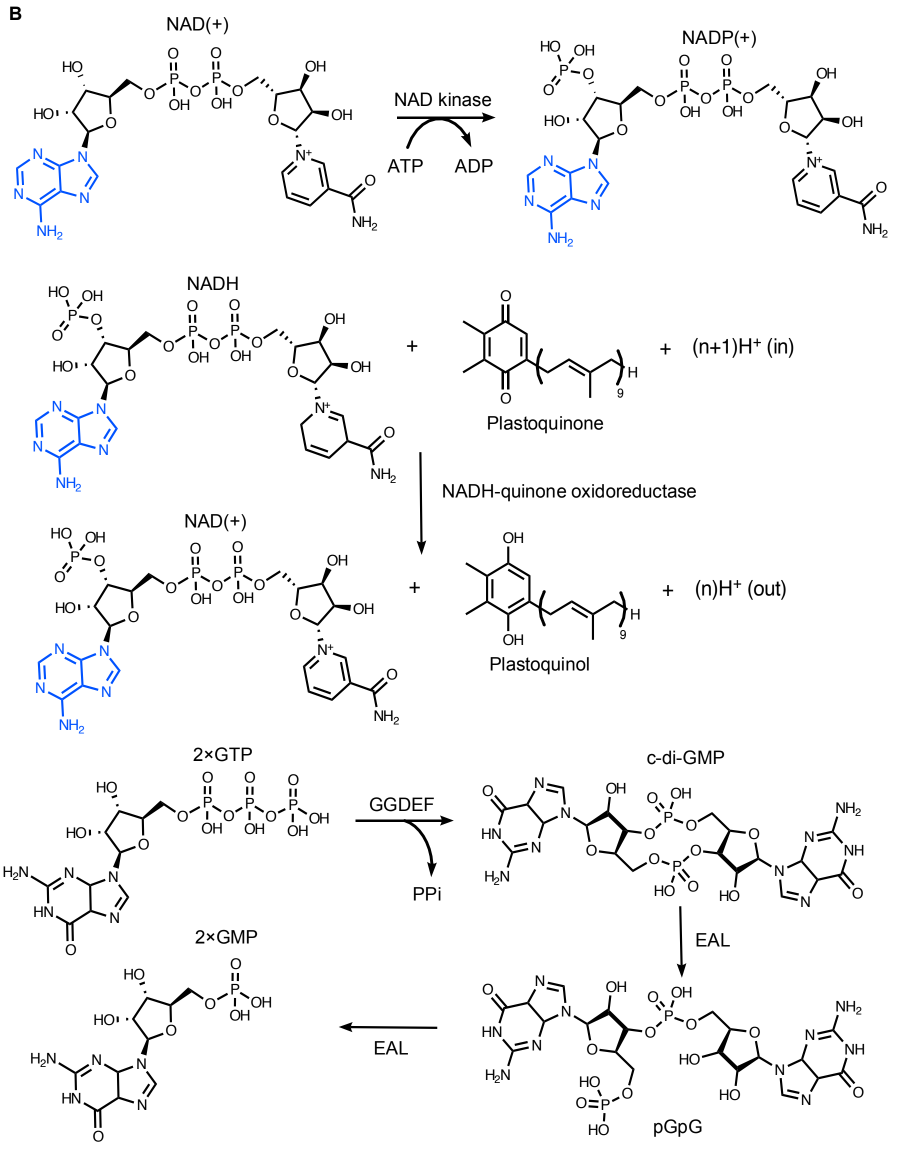


**Supplementary Figure 3.** Genomic contexts of adenine prenyltransferase genes. (A) Comparison of the gene organization surrounding adenine prenyltransferase loci in selected organisms, *Trichormus variabilis* NIES-23, *Anabaena* sp. YBS01, *Agrobacterium fabrum* and *Agrobacterium salinitolerans*. (B) Schematic representation of the putative functions of genes within the *Trichormus variabilis* NIES-23 cluster, including NAD kinase, NADH-quinone oxidoreductase (ndh), and a GGDEF-EAL domain-containing protein.

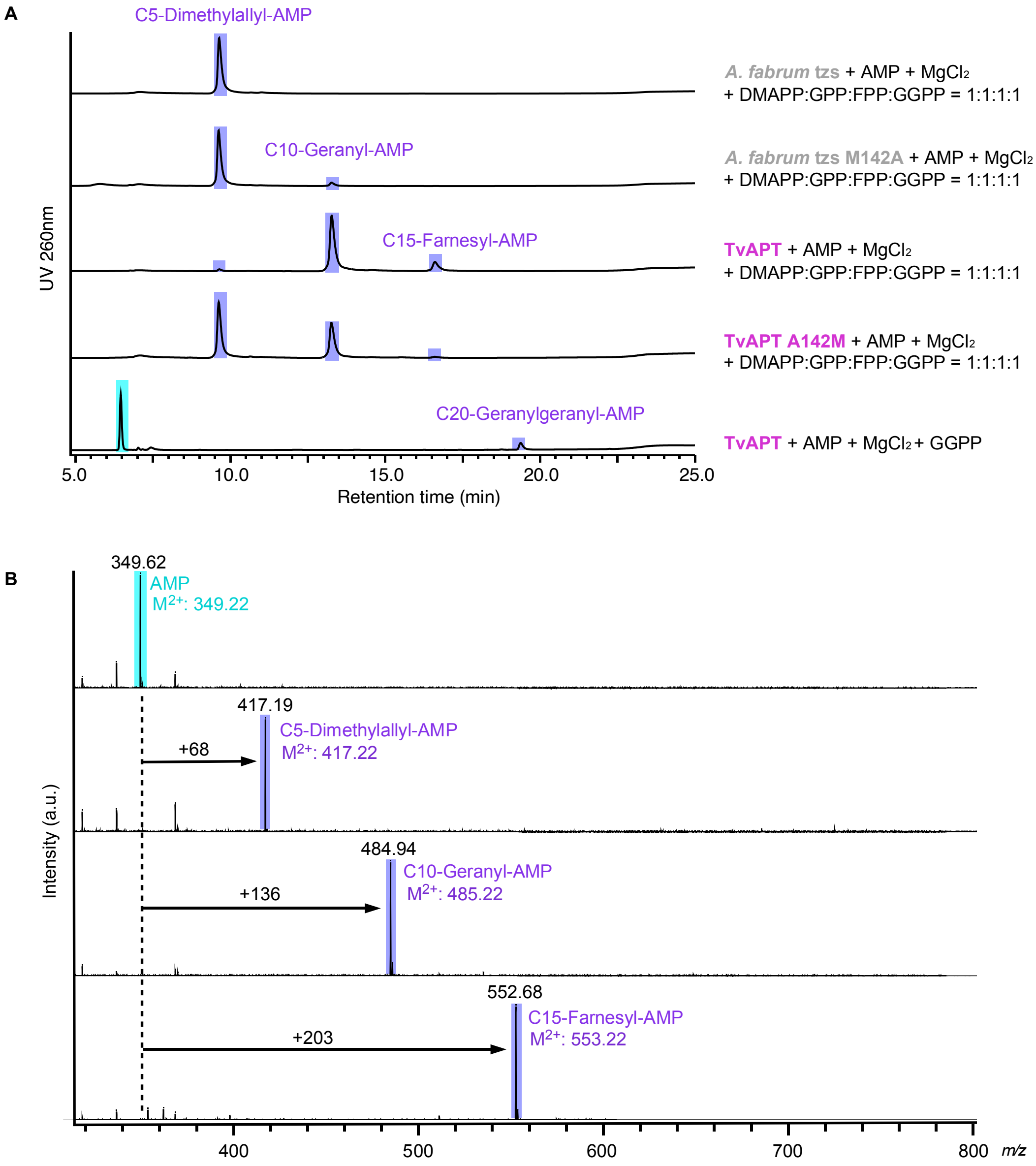


**Supplementary Figure 4.** Comparison of prenyl-donor specificities between TvAPT and tzs from *A. fabrum*. (A) Prenyl-competition assay using C5-C20 prenyl donors. The HPLC peaks corresponding to prenylated AMP were further analyzed by MALDI-TOF MS. (B) MALDI-TOF MS spectra of HPLC-purified prenylated AMP products. The observed mass shifts of +68, +136, and +204 correspond to C5-dimethylallylation, C10-geranylation, and C15-farnesylation, respectively. Peaks corresponding to unmodified AMP and the prenylated products are highlighted in cyan and purple, respectively. No mass peak corresponding to C20-prenylated AMP was detected, likely owing to its low abundance.

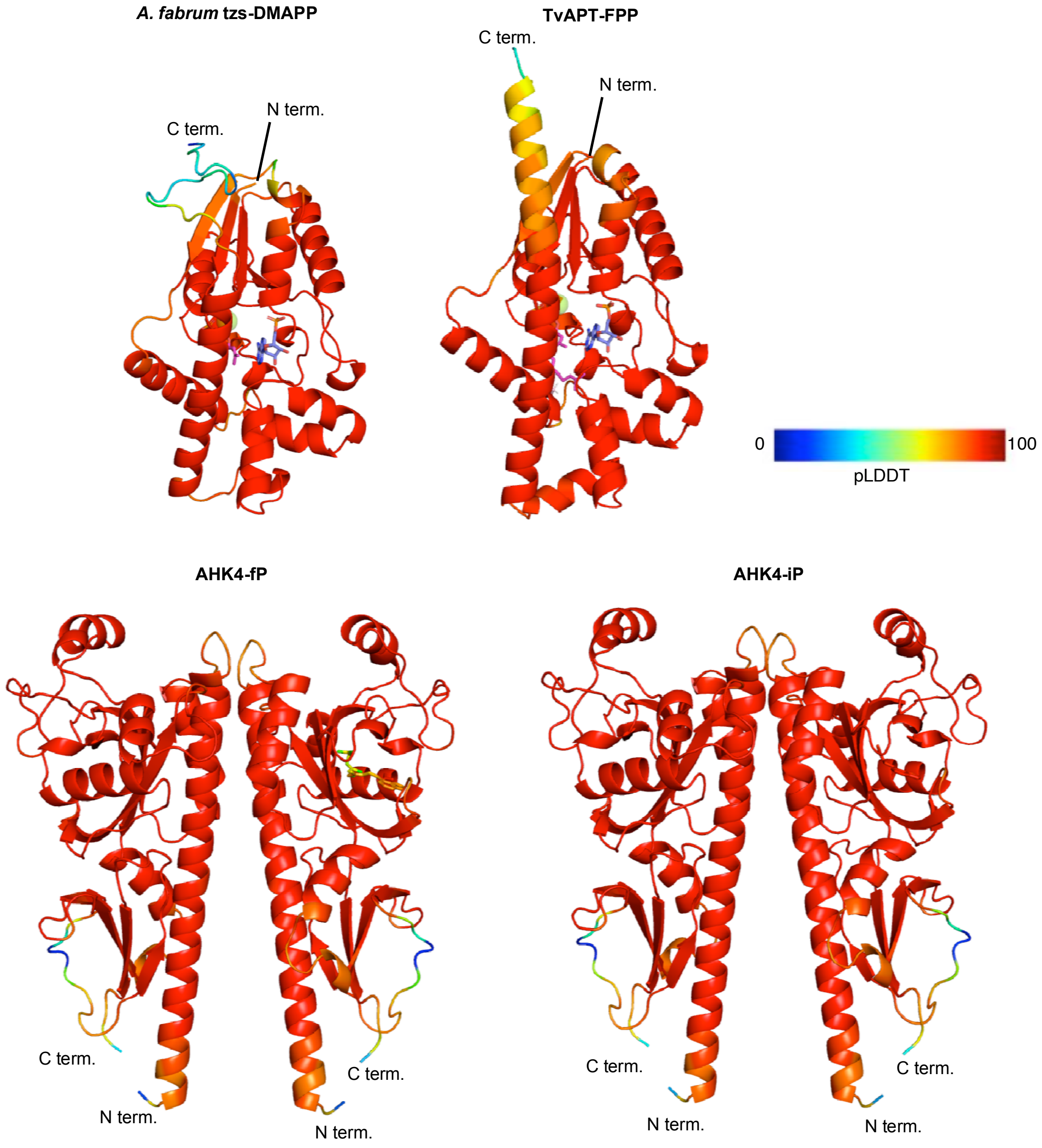


**Supplementary Figure 5.** Predicted structural models of TvAPT, tzs and AHK4. The models were generated using Boltz-2 and are colored according to their predicted local distance difference test (pLDDT) confidence scores. High-confidence regions are indicated in red.

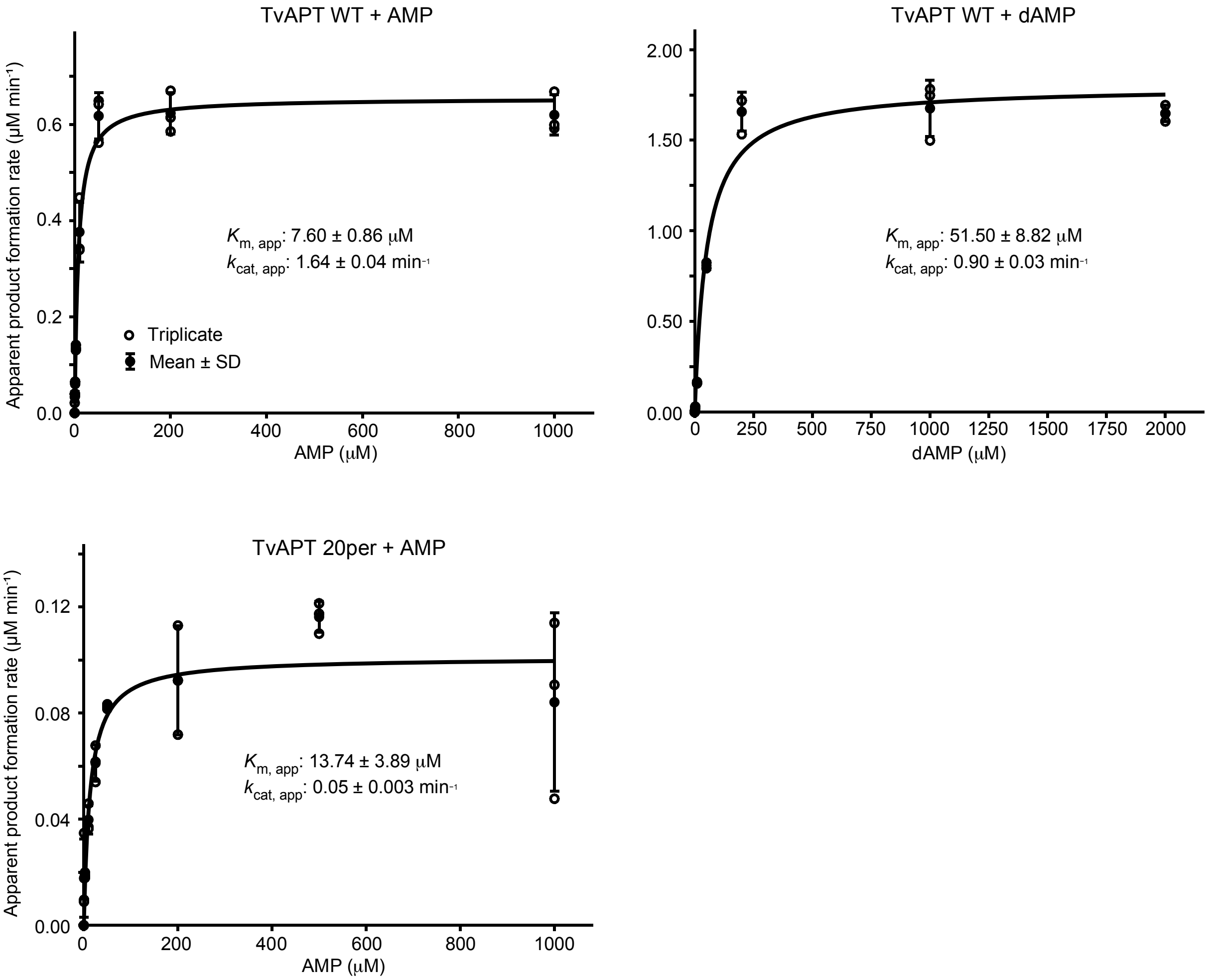


**Supplementary Figure 6.** Steady-state kinetic analysis of TvAPT WT and TvAPT_20per. Initial velocities were determined from the integrated HPLC peak areas of the corresponding prenylated products. The enzyme concentration was 0.4 μM for TvAPT WT with AMP and 2 μM for TvAPT_20per with AMP and TvAPT WT with dAMP. GPP was maintained at a fixed concentration of 500 μM. Data are presented as the mean ± s.d. of three independent experiments. The curves represent fits to the Michaelis–Menten equation.

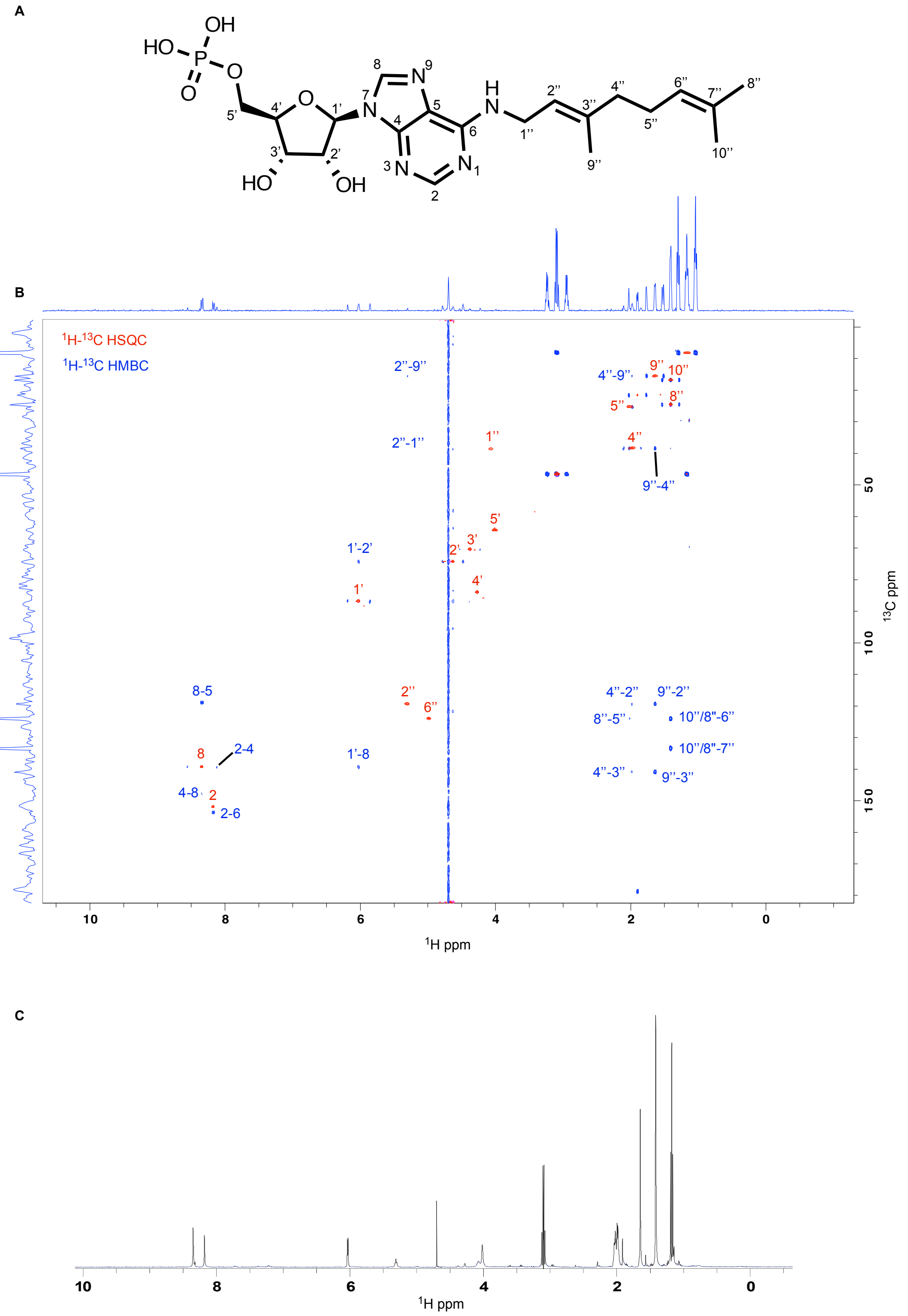


**Supplementary Figure 7.** NMR spectra of N6-geranylated AMP. (A) Chemical structure of N6-geranylated AMP with atom numbering used for the NMR signal assignments. (B) Overlaid $^{1}H$-$^{13}C$ HSQC (red) and $^{1}H$-$^{13}C$ HMBC (blue) spectra, with the corresponding signal assignments indicated. (C) One-dimensional $^{1}H$ NMR spectrum of N6-geranylated AMP recorded in $D_2O$.

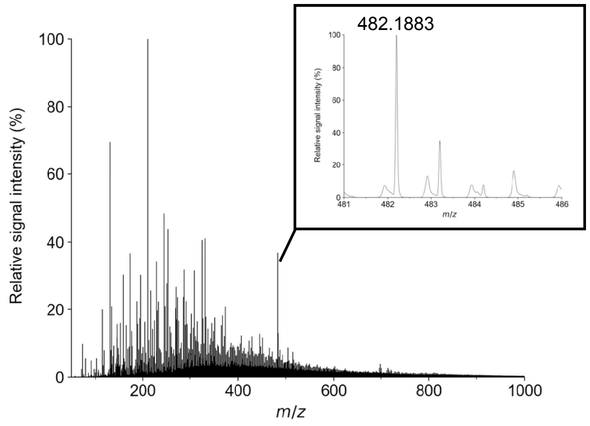


**Supplementary Figure 8.** High-resolution mass spectrometric analysis of N6-geranylated AMP. The peak at *m/z* 482.1883 was assigned to N6-geranylated AMP ($C_{20}H_{29}N_5O_7P^-$; calculated *m/z*, 482.1805).

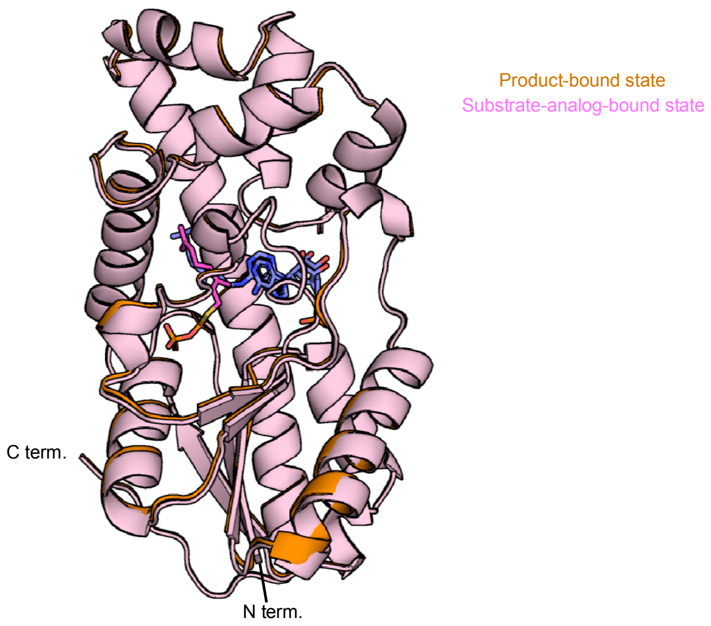


**Supplementary Figure 9.** Structural superposition of the product-bound state (orange) and substrate-analogue-bound state (magenta) of TvAPT_20per.

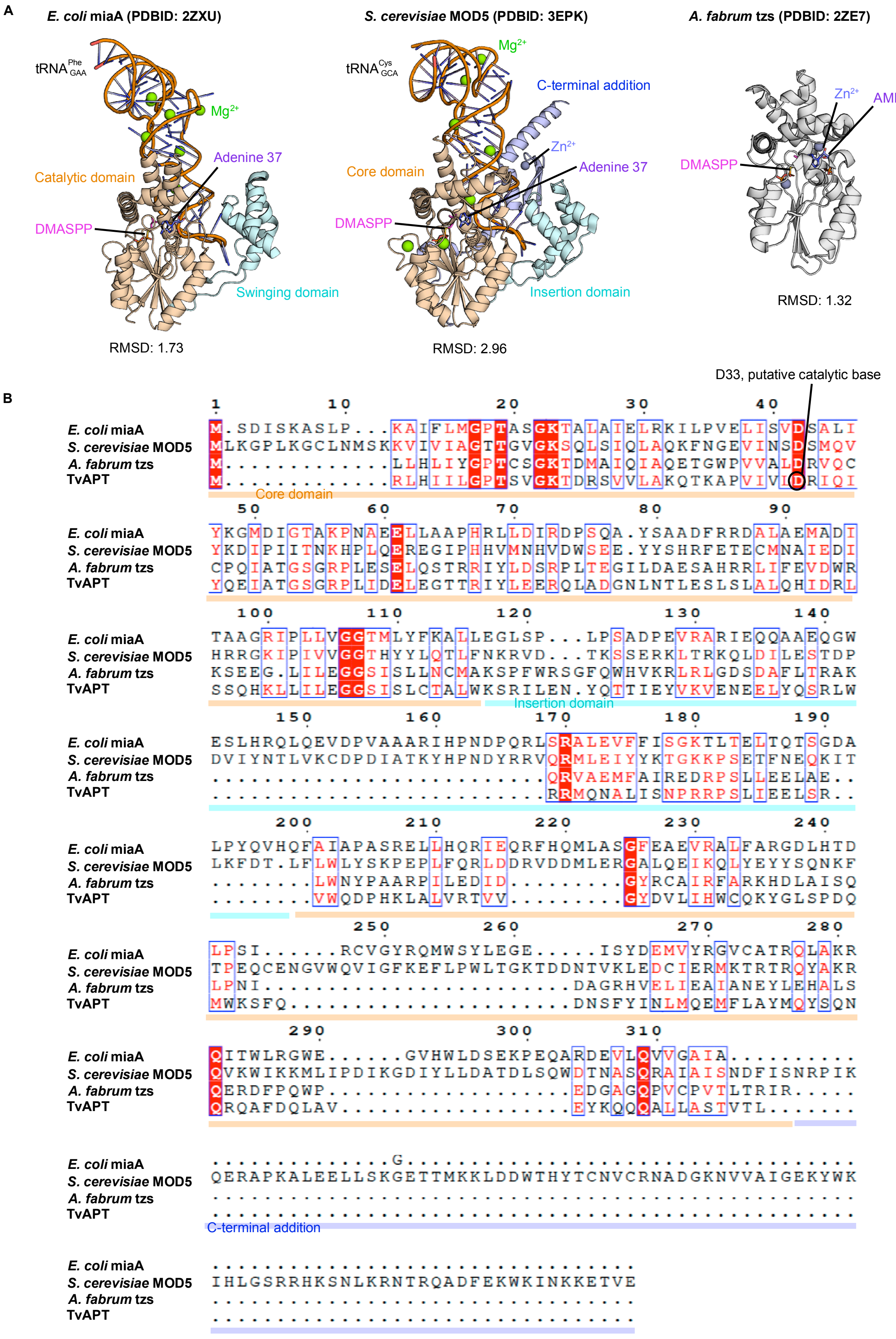
A
E. coli miaA (PDBID: 2ZXU)
S. cerevisiae MOD5 (PDBID: 3EPK)
A. fabrum tzs (PDBID: 2ZE7)
tRNA Phe GAA
Mg2+
Adenine 37
Catalytic domain
DMASPP
Swinging domain
RMSD: 1.73
tRNA Cys GCA
C-terminal addition
Zn2+
Core domain
Insertion domain
RMSD: 2.96
AMP
RMSD: 1.32
B
D33, putative catalytic base
E. coli miaA
S. cerevisiae MOD5
A. fabrum tzs
TvAPT
Core domain
Insertion domain
C-terminal addition

**Supplementary Figure 10.** Structural and sequence comparisons of adenylate prenyltransferases. (A) X-ray crystal structures of MiaA, MOD5, and tzs. RMSD values for Cα-based structural alignments with TvAPT are indicated below each model. (B) Multiple sequence alignment. Regions involved in tRNA binding, the insertion domain, and the extended C-terminal domain are highlighted in cyan and blue.

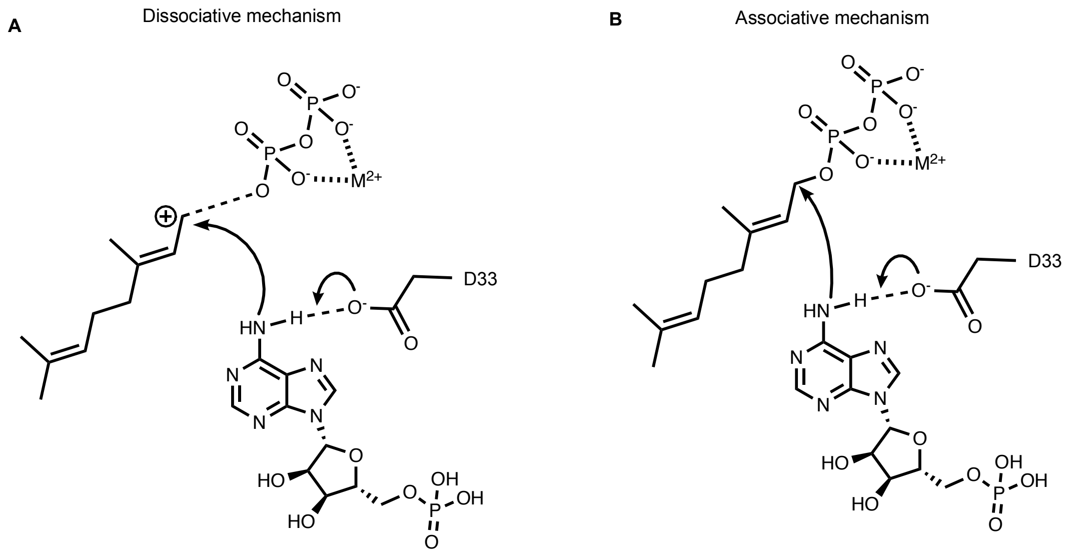


**Supplementary Figure 11.** Putative reaction mechanisms for TvAPT-catalyzed AMP geranylation. (A) An $S_N1$-like stepwise dissociative mechanism involving the departure of pyrophosphate to generate a discrete carbocation intermediate. (B) An $S_N2$-like concerted mechanism involving direct nucleophilic attack on the prenyl donor.

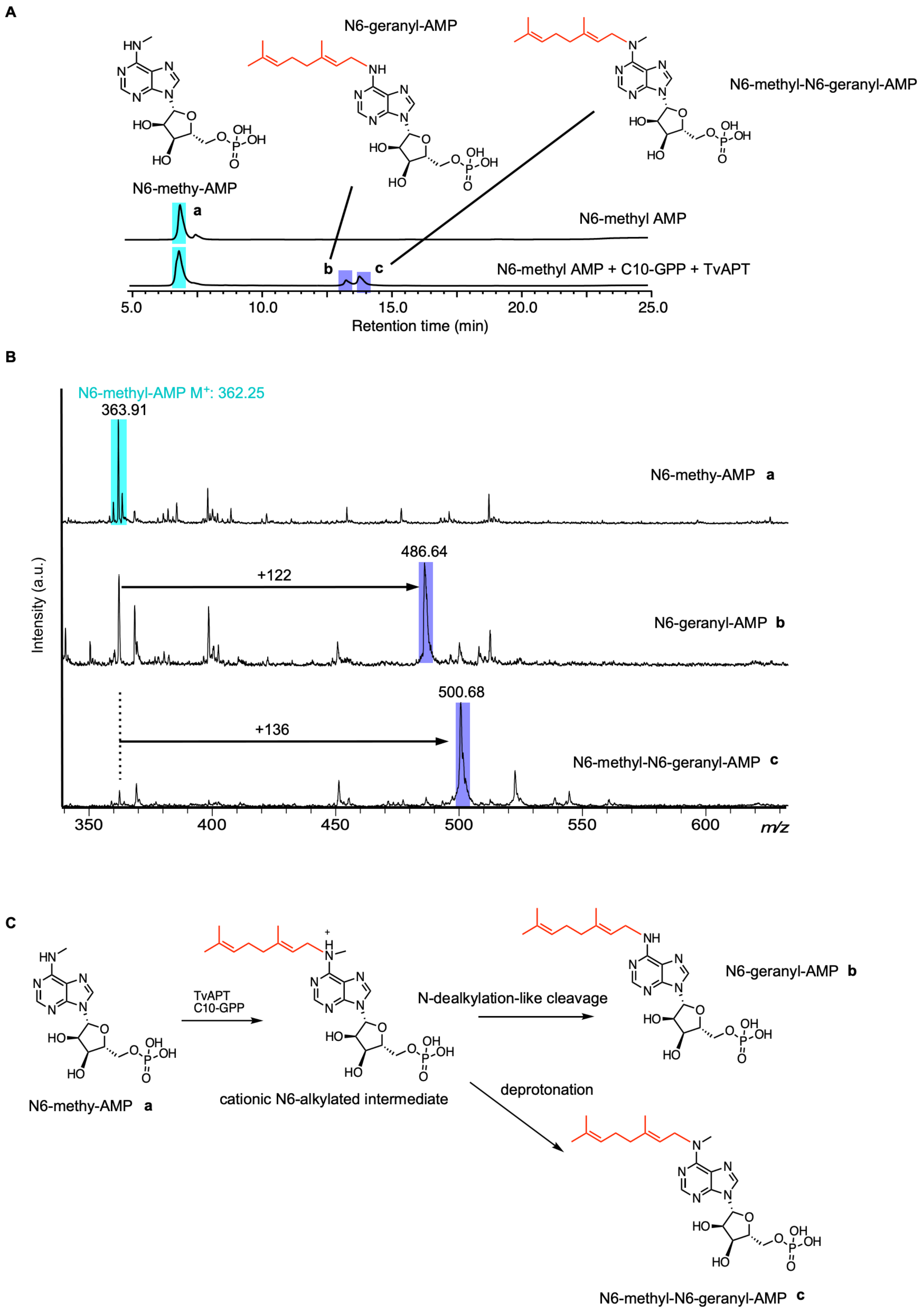
A
N6-geranyl-AMP
N6-methyl-N6-geranyl-AMP
N6-methy-AMP
a
N6-methyl AMP
b
c
N6-methyl AMP + C10-GPP + TvAPT
5.0
10.0
15.0
20.0
25.0
Retention time (min)
B
N6-methyl-AMP M+: 362.25
363.91
N6-methy-AMP a
486.64
+122
N6-geranyl-AMP b
500.68
+136
N6-methyl-N6-geranyl-AMP c
Intensity (a.u.)
350
400
450
500
550
600
m/z
C
N6-methy-AMP a
TvAPT
C10-GPP
cationic N6-alkylated intermediate
N-dealkylation-like cleavage
N6-geranyl-AMP b
deprotonation
N6-methyl-N6-geranyl-AMP c

**Supplementary Figure 12.** TvAPT-catalyzed geranylation of N6-methyl-AMP and the proposed mechanism involving a cationic N6-alkylated intermediate. (A) HPLC chromatogram of the TvAPT-catalyzed reaction with N6-methyl-AMP. Peaks **b** and **c** were collected and analyzed by MALDI-TOF MS. (B) MALDI-TOF mass spectra of peaks **a**–**c**. Mass shifts of +122 and +136 Da relative to N6-methyl-AMP were assigned to geranylation with and without demethylation, respectively. (C) Proposed reaction mechanism involving a cationic N6-methyl-N6-geranyl intermediate. N-Dealkylation-like cleavage of the N6–CH3 bond would yield N6-geranyl-AMP, whereas deprotonation would yield N6-methyl-N6-geranyl-AMP.

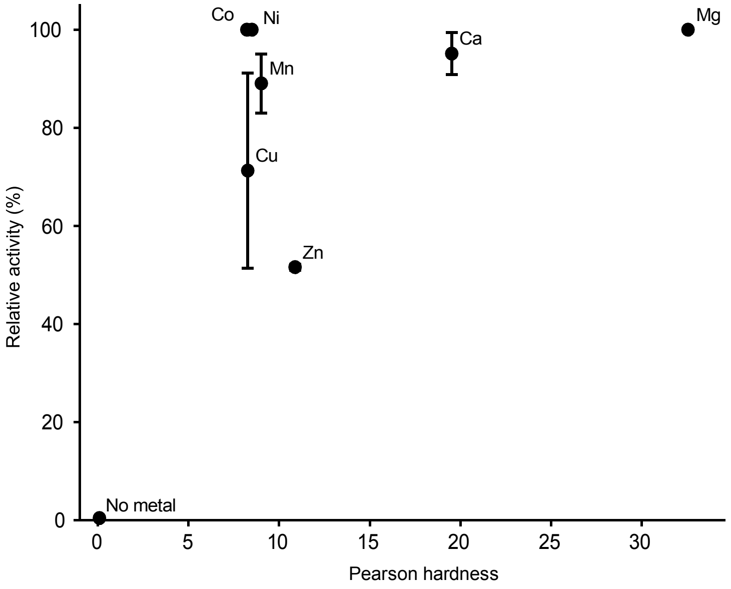


**Supplementary Figure 13.** Metal dependence of TvAPT-catalyzed AMP geranylation. Relative activities were calculated from the HPLC peak areas of the geranylated product, normalized to the activity in the presence of $Mg^{2+}$, which was set to 100%, and plotted against the Pearson hardness of each metal ion. Values are presented as the mean ± s.d. of three independent experiments.

**A**

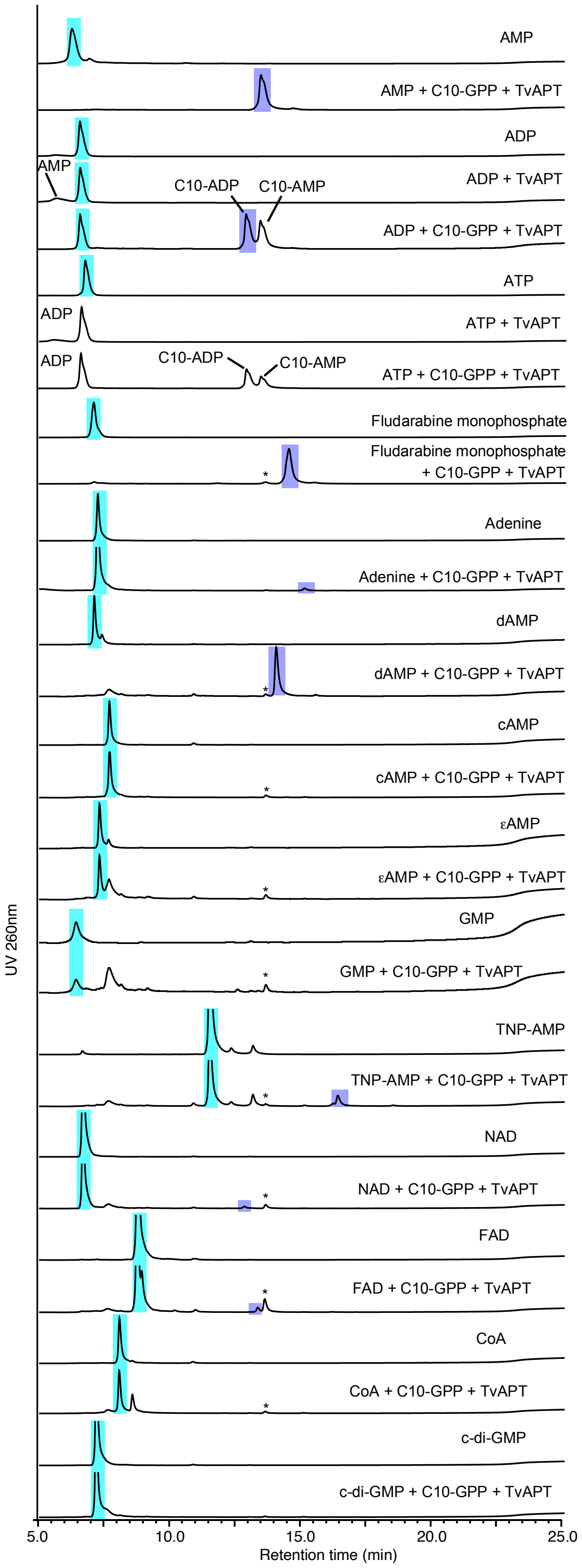
AMP
AMP + C10-GPP + TvAPT
ADP
AMP
ADP + TvAPT
C10-ADP
C10-AMP
ADP + C10-GPP + TvAPT
ATP
ADP
ATP + TvAPT
ADP
C10-ADP
C10-AMP
ATP + C10-GPP + TvAPT
Fludarabine monophosphate
Fludarabine monophosphate + C10-GPP + TvAPT
Adenine
Adenine + C10-GPP + TvAPT
dAMP
dAMP + C10-GPP + TvAPT
cAMP
cAMP + C10-GPP + TvAPT
εAMP
εAMP + C10-GPP + TvAPT
GMP
GMP + C10-GPP + TvAPT
TNP-AMP
TNP-AMP + C10-GPP + TvAPT
NAD
NAD + C10-GPP + TvAPT
FAD
FAD + C10-GPP + TvAPT
CoA
CoA + C10-GPP + TvAPT
c-di-GMP
c-di-GMP + C10-GPP + TvAPT
UV 260nm
5.0
10.0
15.0
20.0
25.0
Retention time (min)

B

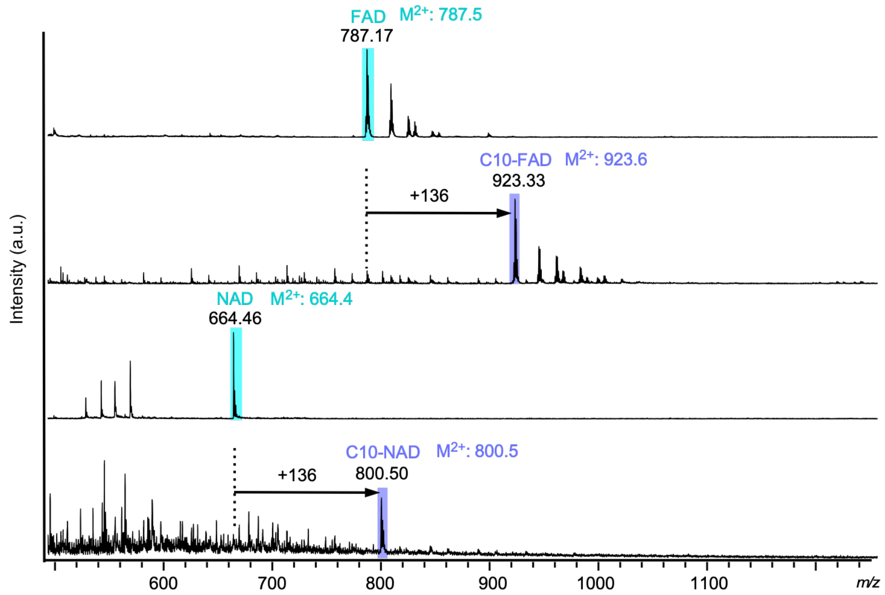


**Supplementary Figure 14.** *In vitro* geranylation of diverse adenine-containing substrates. (A) HPLC chromatograms of reaction mixtures containing the indicated adenine derivatives. Peaks corresponding to unmodified substrates and geranylated products are highlighted in cyan and purple, respectively. Asterisks indicate peaks assigned to AMP-derived products, generated by the geranylation of endogenous AMP co-purified with TvAPT. (B) MALDI-TOF MS spectra of purified geranylated NAD and FAD.

A

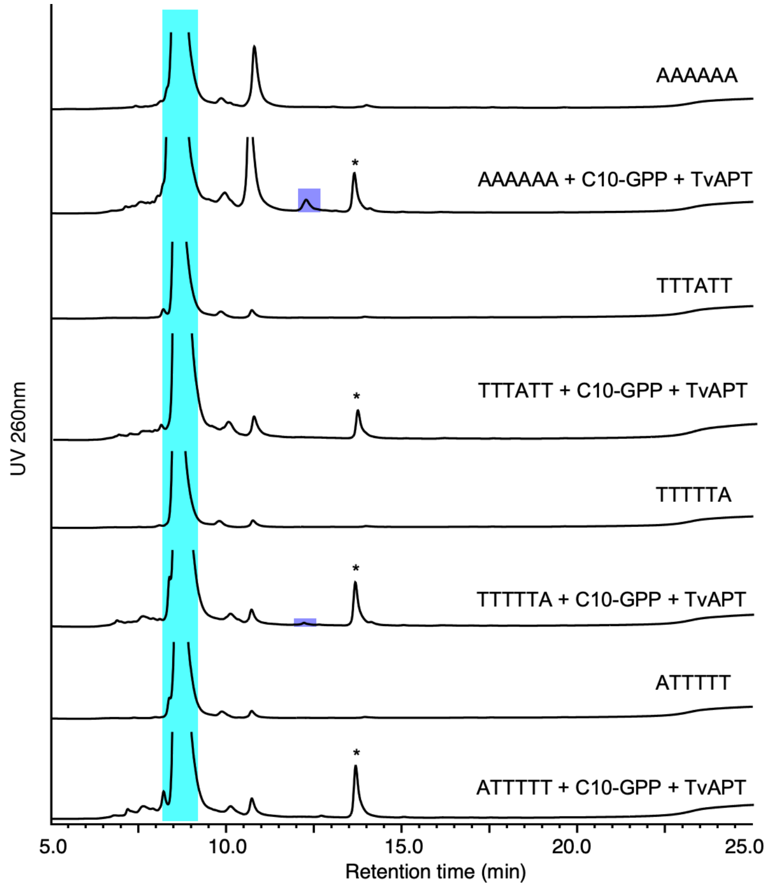
AAAAAA
*
AAAAAA + C10-GPP + TvAPT
TTTATT
*
TTTATT + C10-GPP + TvAPT
TTTTTA
*
TTTTTA + C10-GPP + TvAPT
ATTTTT
*
ATTTTT + C10-GPP + TvAPT
UV 260nm
5.0
10.0
15.0
20.0
25.0
Retention time (min)

B

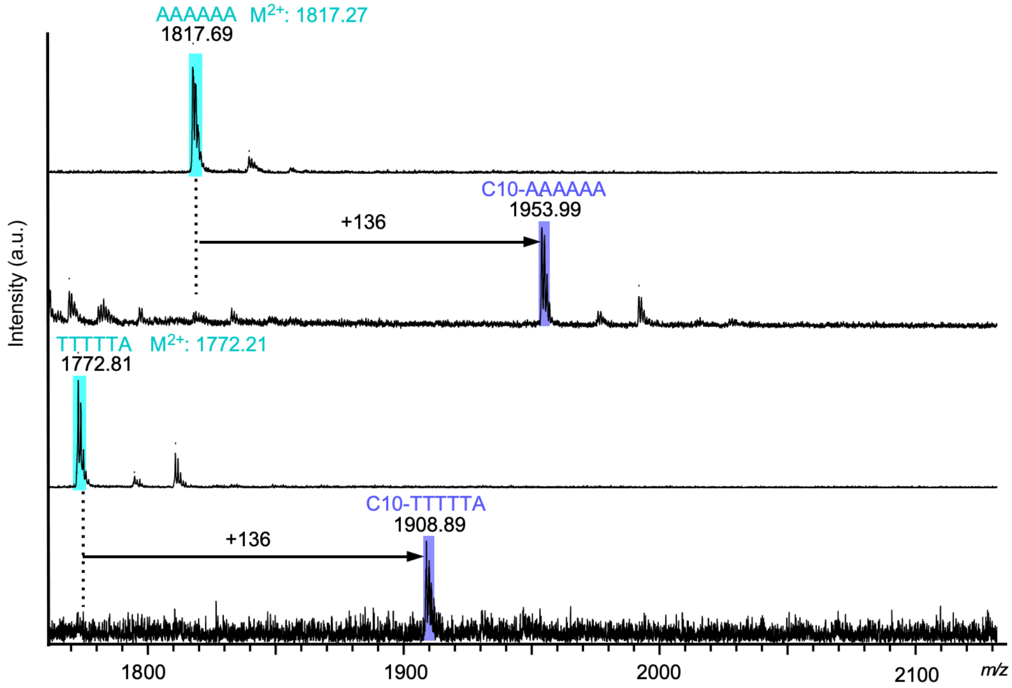
AAAAAA M2+: 1817.27
1817.69
C10-AAAAAA
1953.99
+136
TTTTTA M2+: 1772.21
1772.81
C10-TTTTTA
1908.89
+136
Intensity (a.u.)
1800
1900
2000
2100
m/z

**Supplementary Figure 15.** *In vitro* geranylation of 6-mer single-stranded oligodeoxynucleotides. (A) HPLC chromatograms of reaction mixtures containing the indicated oligonucleotides. Peaks corresponding to unmodified substrates and geranylated products are highlighted in cyan and purple, respectively. Asterisks indicate peaks assigned to AMP-derived products, generated by the geranylation of endogenous AMP co-purified with TvAPT. (B) MALDI-TOF MS spectra of purified geranylated AAAAAA and TTTTTA.

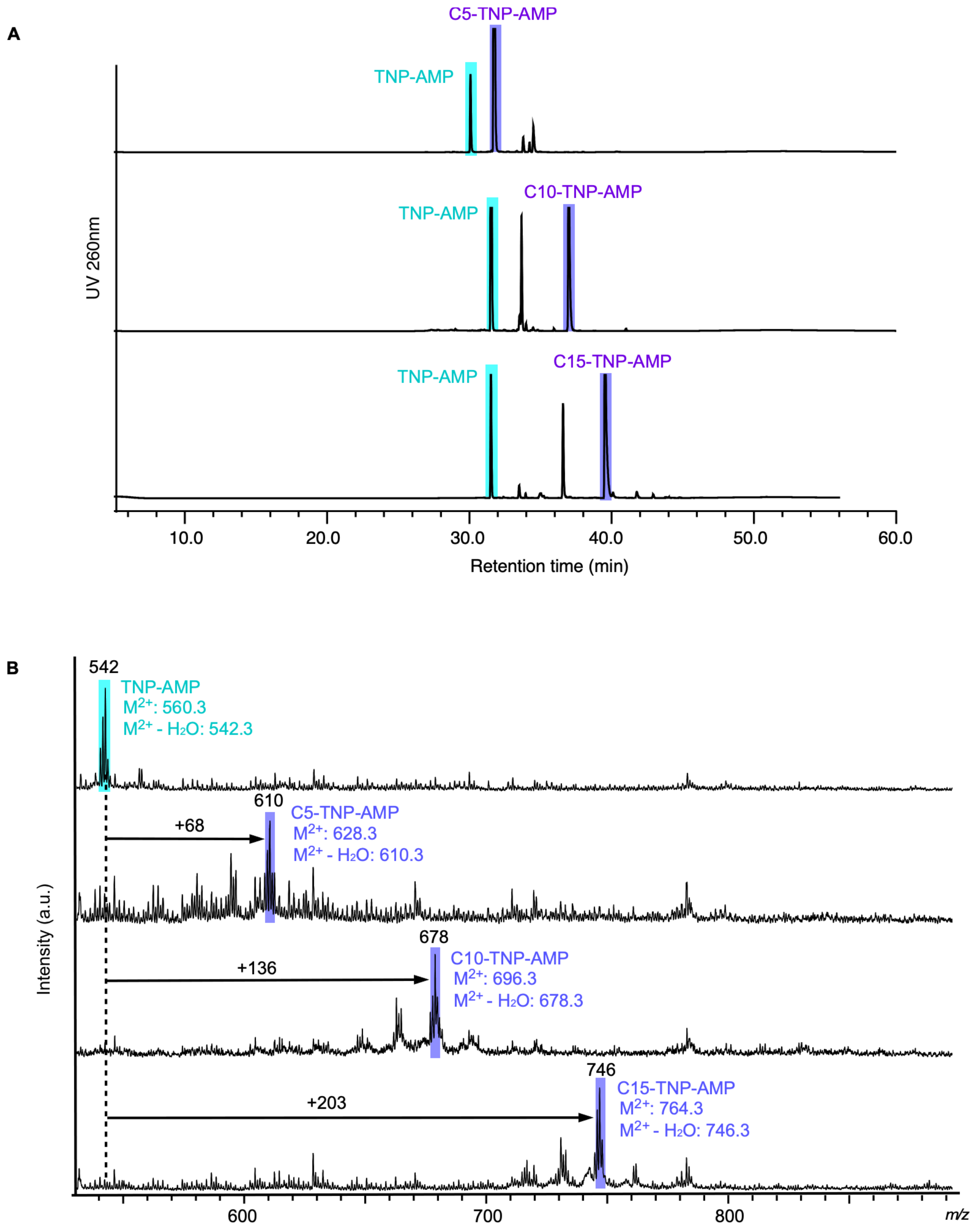


**Supplementary Figure 16.** Preparation of prenylated TNP-AMP derivatives. (A) Preparative HPLC chromatograms of TNP-AMP enzymatically prenylated by TvAPT. (B) MALDI-TOF MS spectra of the purified products. In both panels, peaks corresponding to unmodified TNP-AMP and the prenylated products are highlighted in cyan and purple, respectively. The prenylated products were detected in a dehydrated form.

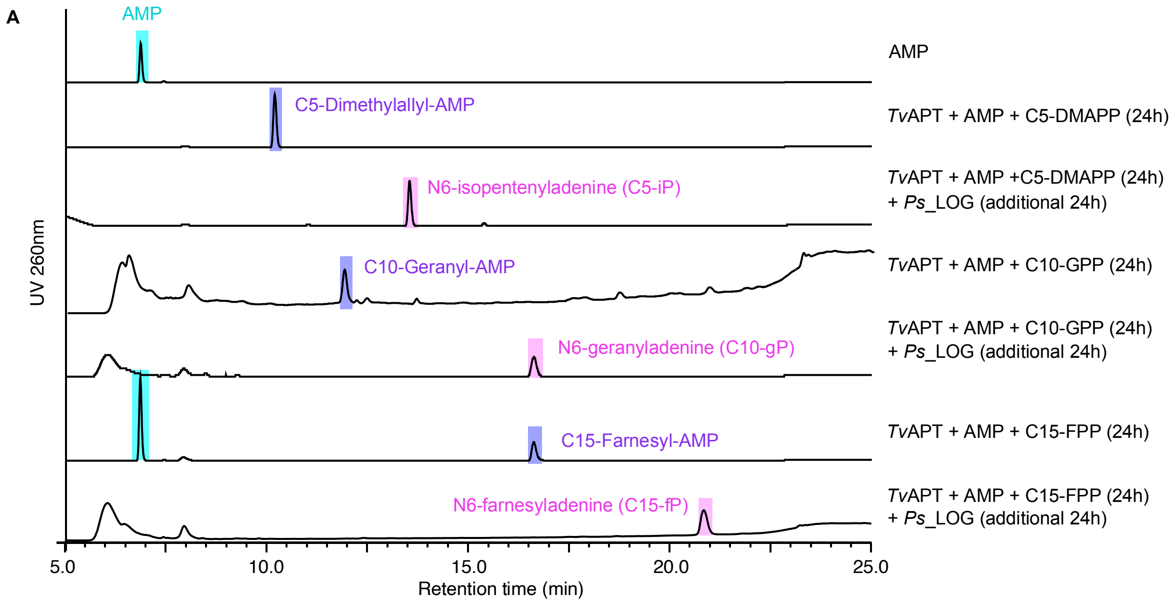


**Supplementary Figure 17.** HPLC chromatograms of the reaction mixtures used for the enzymatic synthesis of cytokinin analogues. Peaks corresponding to unmodified AMP, prenylated AMP, and deribosylated prenylated adenines are highlighted in cyan, purple, and magenta, respectively.

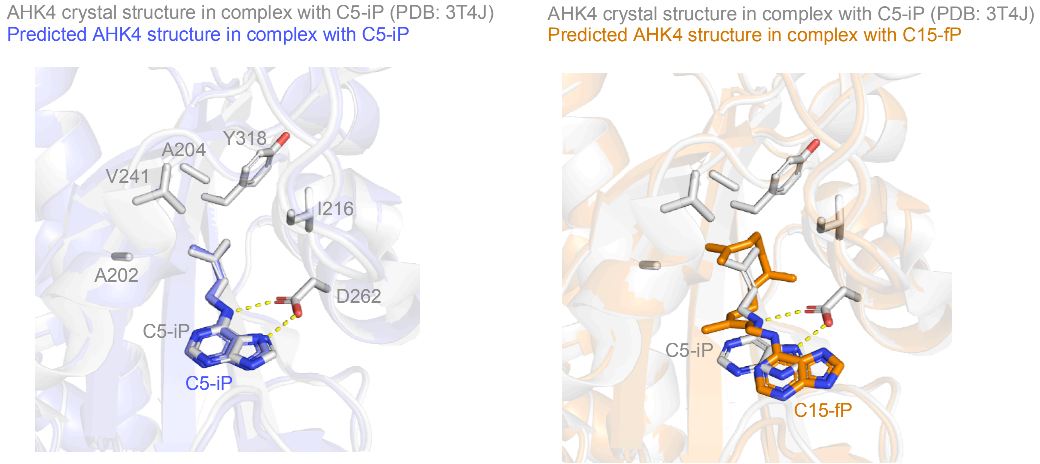


**Supplementary Figure 18.** Predicted interactions of C5-iP (blue) and C15-fP (orange) with AHK4. The X-ray crystal structure of AHK4 in complex with C5-iP (gray) was superimposed on the predicted AHK4 structures in complex with C5-iP and C15-fP. Prenyl-recognition residues and cytokinin ligands are shown as stick models.

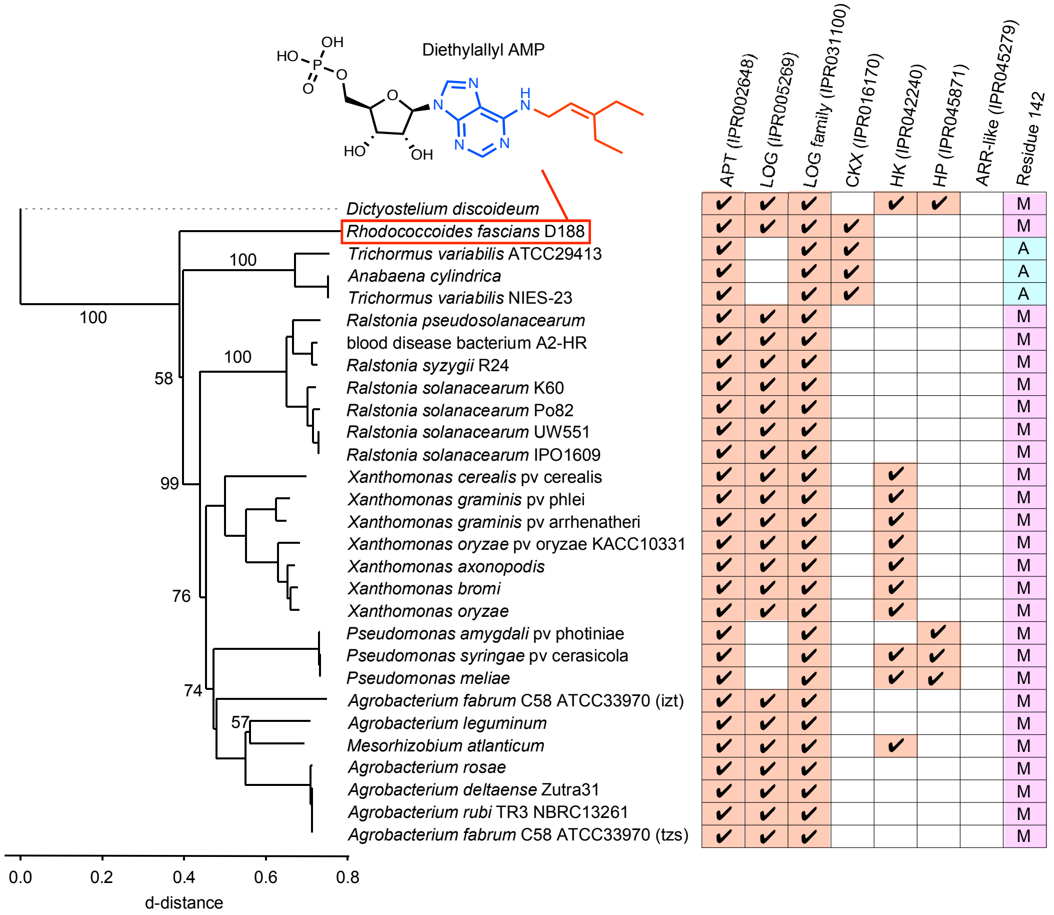


**Supplementary Figure 19.** Phylogenetic analysis of bacterial adenylate prenyltransferases putatively involved in cytokinin biosynthesis. Bootstrap values are indicated at the nodes. Check marks denote the presence of genes encoding APT, LOG, LOG-like proteins, CKX, HK, and ARR-like proteins within the corresponding genomes. The amino acid residue at position 142 (A, alanine; M, methionine) is shown in the rightmost column. The chemical structure of the N6-diethylallylated AMP produced by the *Rhodococcus fascians* AIPT is depicted above the phylogenetic tree. The proteins included in this analysis correspond to the cluster circled in orange in the sequence similarity network shown in Supplementary Fig. 2.

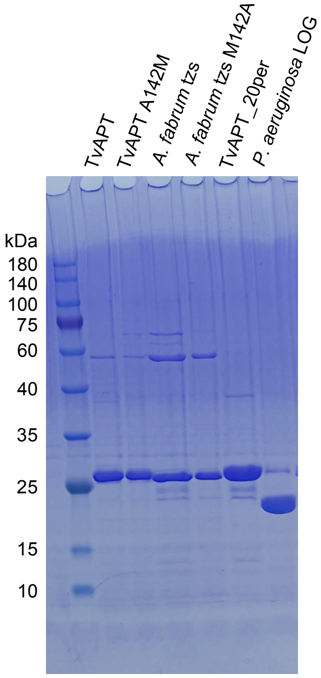


**Supplementary Figure 20.** SDS-PAGE analysis of His-tag-purified proteins. The ExcelBand 3-color Regular Range Protein Marker (9–180 kDa; SMOBIO) was used as the molecular-weight marker.

**Supplementary Table 1. Predicted binding constants of cytokinin derivatives with AHK4.**

| Cytokinin | Apparent $K_D$ (nM) reported in Ref. 49 | Apparent $K_D$ (nM) reported in Ref. 50 | Predicted binding scores (kcal/mol) by Boltz-2 |
|---|---|---|---|
| *trans*-Zeatin | 3.9 | 4.4 ± 1.1 | -8.3 |
| *cis*-Zeatin | 830 | 454 ± 134 | -8.4 |
| 6BP | 300 | 64 ± 1.5 | -6.3 |
| C5-iP | 17 | 3.5 ± 1.6 | -7.6 |
| C10-gP | - | - | -8.4 |
| C15-fP | - | - | -7.4 |
| Thidiazuron | 40 | - | -8.7 |

**Supplementary Table 2. List of primers.**

| Name | Sequence |
|---|---|
| AIPT_M142A_f | TGTGGCCGAGGCGTTCGCCATCCG |
| AIPT_M142A_r | CGCTGCTTTGCGCGC |
| TvAPT_A142M_f | TATGCAAAACATGCTTATCAGCAATCCGC |
| TvAPT_A142M_r | CGACGCCATAAGCGG |

**Supplementary Table 3. List of amino acid sequences.**

| Protein | Amino acid sequence |
|---|---|
| TvAPT | MGSSHHHHHHSSGLVPRGSHMASMRLHIILGPTSVGKTDRSVVL<br>AKQTKAPVIVLDRIQIYQEIATGSGRPLIDELEGTTRIYLEERQLA<br>DGNLNTLESLSLALQHIDRLSSQHKLLILEGGSISLCTALWKSRIL<br>ENYQTTIEYVKVENEELYQSRLWRRMQNALISNPRRPSLIEELSR<br>VWQDPHKLALVRTVVGYDVLIHWCQKYGLSPDQMWKSFQDNS<br>FYINLMQEMFLAYMQYSQNQRQAFDQLAVEYKQQQALLASTVT<br>L |
| TvAPT_20per | MGSSHHHHHHSSGLVPRGSHMRLHIILGPTSVGKTKRSVKLAKQ<br>TKAPVIVLDRIQIYQEIATGSGRPLPDELEGTKRIYLTERSLADGN<br>LNTDEALSLALQHIDRLSSQHKLLILEGGSISLCTALWKSRILENY<br>QTTIEYVQVENEELYQSRLERRLQNALQSNPRRPSLIEELSRVWQ<br>DPHRLALVRTVVGYDVLIQWCQKLGLSPDQLIKQFQDLSFESNL<br>LQEMFLAYRQLSQNQRQAFDQLAVEYKQQQALLASTLTL |
| *A. fabrum* tzs | MGSSHHHHHHSSGLVPRGSHMASMLLHLIYGPTCSGKTDMAIQI<br>AQETGWPVVALDRVQCCPQIATGSGRPLESELQSTRRIYLDSRPL<br>TEGILDAESAHRRLIFEVDWRKSEEGLILEGGSISLLNCMAKSPF<br>WRSGFQWHVKRLRLGDSDAFLTRAKQRVAEMFAIREDRPSLLEE<br>LAELWNYPAARPILEDIDGYRCAIRFARKHDLAISQLPNIDAGRH<br>VELIEAIANEYLEHALSQERDFPQWPEDGAGQPVCPVTLTRIR |
| *P. aeruginosa* LOG | MGMTLRSVCVFCGASPGASPVYQEAAVALGRHLAERGLTLVYG<br>GGAVGLMGTVADAALAAGGEVIGIIPQSLQEAEIGHKGLTRLEV<br>VDGMHARKARMAELADAFIALPGGLGTLEELFEVWTWGQLGY<br>HAKPLGLLEVNGFYDPLLTFLDHLVDERFVRAEHRGMLQRGASP<br>EALLDALAAWTPSVAPKWVDRTPQLEHHHHHH |
| *A. thaliana* AHK4 (149-418) | MDDANKIRREEVLVSMCDQRARMLQDQFSVSVNHVHALAILVS<br>TFHYHKNPSAIDQETFAEYTARTAFERPLLSGVAYAEKVVNFERE<br>MFERQHNWVIKTMDRGEPSPVRDEYAPVIFSQDSVSYLESLDM<br>MSGEEDRENILRARETGKAVLTSPFRLLETHHLGVVLTFPVYKSS<br>LPENPTVEERIAATAGYLGGAFDVESLVENLLGQLAGNQAIVVH<br>VYDITNASDPLVMYGNQDEEADRSLSHESKLDFGDPFRKHKMIC<br>RYHQKA |

**Supplementary Table 4. Data collection and refinement statistics for TvAPT_20per.**

| | TvAPT_20per (geranyl-AMP) | TvAPT_20per (AMP and GSPP) |
|---|---|---|
| Space group | C121 | C121 |
| Unit cell parameters | | |
| a (Å) | 128.5 | 128.2 |
| b (Å) | 38.2 | 38.8 |
| c (Å) | 54.7 | 54.9 |
| α (degree) | 90.0 | 90.0 |
| β (degree) | 94.5 | 94.2 |
| γ (degree) | 90.0 | 90.0 |
| Wavelength (Å) | 1.00 | 1.00 |
| Resolution (Å) | 43.2-1.70 (1.73-1.70) | 43.2-1.85 (1.89-1.85) |
| No. of reflections | 192,918 | 154,190 |
| No. of unique reflections | 29,478 | 23.247 |
| Completeness (%) | 99.9 (99.9) | 99.9 (99.8) |
| I/sig (I) | 24.6 (2.0) | 19.3 (2.5) |
| $R_{merge}$ | 0.033 (0.867) | 0.062 (0.862) |
| $CC_{1/2}$ | 1.000 (0.798) | 0.999 (0.851) |
| *B* of Wilson plot (Å$^2$) | 26.7 | 23.5 |
| $R_{work}$ | 0.212 | 0.183 |
| $R_{free}^{a}$ | 0.247 | 0.224 |
| RMSD of geometry | | |
| Bond length (Å) | 0.01 | 0.01 |
| Bond angles (deg) | 1.75 | 1.74 |
| Geometry | | |
| Ramachandran outlier (%) | 0.4 | 0.0 |
| Ramachandran favored (%) | 99.6 | 100.0 |
| PDB entry | 24WM | 27DP |

[a.] $R_{free}$ is the *R*-factor calculated using 5% of the reflections randomly selected and omitted from refinement.

**Supplementary Table 5. Chemical shift assignments for geranylated AMP.**

| Atom | $^{1}H$ (ppm), multiplicity | $^{13}C$ (ppm) |
|---|---|---|
| 2 | 8.17, s | 151.9 |
| 4 | | 148.1 |
| 5 | | 118.9 |
| 6 | | 154 |
| 8 | 8.34, s | 139.3 |
| 1' | 6.03, d (6.0 Hz) | 86.8 |
| 2' | 4.62 | 74.2 |
| 3' | 4.38, t (5.3 Hz) | 70.4 |
| 4' | 4.27, br | 83.9 |
| 5' | 4.01, br | 64.3 |
| 1" | 4.06, br | 38.6 |
| 2" | 5.3, t (7.0 Hz) | 119.2 |
| 3" | | 140.8 |
| 4" | 1.98, d (6.0 Hz) | 38.3 |
| 5" | 2.01, t (6.8 Hz) | 25.2 |
| 6" | 4.99, t (7.2 Hz) | 123.9 |
| 7" | | 133.4 |
| 8" | 1.41, s | 24.6 |
| 9" | 1.64, s | 15.5 |
| 10" | 1.15 | 18.1 |
